# The usage of human IGHJ genes follows a particular nonrandom selection: The recombination signal sequence affects the usage of human IGHJ genes

**DOI:** 10.1101/792085

**Authors:** Bin Shi, Xiaoheng Dong, Qingqing Ma, Suhong Sun, Long Ma, Jiang Yu, Xiaomei Wang, Juan Pan, Xiaoyan He, Danhua Su, Xinsheng Yao

**Affiliations:** Department of Immunology, Center of Immunomolecular Engineering, Innovation & Practice Base for Graduate Students Education, Zunyi Medical University, Zunyi, China; Departmentof Breast Surgery, The first Affiliated Hospital of ZunYi Medical University, Zunyi, China; Department of Laboratory Medicine, Affiliated Hospital of Zunyi Medical University, Zunyi, China; School of Laboratory Medicine, Zunyi Medical University, Zunyi, China

## Abstract

The formation of the B cell receptor (BCR) heavy chain variable region is derived from the germline V(D)J gene rearrangement according to the “12/23” rule and the “beyond 12/23” rule. The usage frequency of each V(D)J gene in the peripheral BCR repertoires is related to the initial recombination, self-tolerance selection, and the clonal proliferative response. However, their specific differences and possible mechanisms are still unknown. We analyzed in-frame and out-of-frame BCR-H repertoires from human samples with physiological and various pathological conditions by high-throughput sequencing. Our results showed that IGHJ gene frequency follows a similar pattern where IGHJ4 is used at high frequency (>40%), IGHJ6/IGHJ3/IGHJ5 is used at medium frequencies (10%∼20%), and IGH2/IGHJ1 is used at low frequency (<4%) under whether physiological or various pathological conditions. Furthermore, analysis of the recombination signal sequences suggested that the conserved nonamer and heptamer and certain 23 bp spacer length may affect the initial IGHD-IGHJ recombination, which results in different frequencies of IGHJ genes among the initial BCR-H repertoire. Based on this “background repertoire”, we recommend that re-evaluation and further investigation are needed when analyzing the significance and mechanism of IGHJ gene frequency in self-tolerance selection and the clonal proliferative response.

## INTRODUCTION

The diversity of the initial vertebrate B cell receptor (BCR) originates from the recombination of multiple germline genes (V(D)J) and insertion and deletion during the recombination process. There is a consensus recombination signal sequence (RSS) (1) at the 5’ or 3’ end of each V(D)J gene segment that participates in recombination according to the “12/23” rule (2, 3, 4) and the “beyond 12/23” rule (5). In addition, recombination-activating gene (RAG) enzymes, terminal deoxynucleotidyl transferase (TDT), heterodimer-KU70/KU80, DNA-dependent protein kinase (DNA-PK/Artemis), DNA ligase IV (XRCC4) and other proteins are involved in the complex V(D)J recombination process (4, 6).

Theoretically, the usage frequency of V(D)J gene segments is random in the pro-B cell or pre-B cell recombination process (before autoantigen selection). However, in vitro experiments in B cell lines confirmed that V(D)J gene segments contribute unequally to the primary repertoire, and the consensus heptamer and nonamer sequences of the RSSs are considered the major factor (7). The contributing factors may relate to the usage frequency of V(D)J gene segments. The usage of proximal and distal gene segments in recombination is not random; for example, the JH-proximal VH gene of pre-B cell lines has a preferential usage (8), and VH near Cu may be preferred during early rearrangement (9). During pre-B cell differentiation and development, the initial DH-JH rearrangements employ more 3’ (JH-proximal) DH segments (10); however, Feeney AJ et al found that there is no apparent preference for the more JK-proximal over the more JK-distal genes in the proximal region (11). In addition, compared with RSSs with one or more base mutations, the corresponding gene subfamily of RSSs with a consensus heptamer/nonamer (conserved) has preferred usage (3, 12, 13, 14, 15). Moreover, the usage frequency of the corresponding gene segment will be affected when the lengths of the 12 bp spacer/23 bp spacer in RSSs increase or decrease (12, 13, 15) and when the base sequences of the 12 bp spacer/23 bp spacer in RSS change (16,17,18).

However, these results are derived from experiments based on B cell lines in vitro, and whether RSSs influence the V(D)J usage frequency of initial repertoires in vivo is unclear. The difference in each V(D)J usage frequency in the peripheral B cell repertoires is mainly derived from the selection of self-tolerance and the response of clonal proliferation (8, 19, 20, 21). How the difference in usage frequency of each V(D)J gene segment in initial repertoires influences the peripheral repertoire has not been clarified and has received little attention.

With the development of high-throughput sequencing (HTS) analysis for V(D)J tracking, analyzing each V(D)J usage frequency in individual BCR-H repertoires is now possible. We have broadly analyzed the composition characteristics of BCR-H repertoires by HTS since 2013 and found that the human IGHJ4 gene subfamily has the highest usage frequency in physiological and various pathological conditions, followed by IGHJ6, IGHJ3 and IGHJ5 with medium usage frequency and by IGHJ1 and IGHJ2 with significantly low usage frequency. Additionally, the usage frequency of 6 IGHJ gene families shows amazing consistency by analyzing the BCR-H sequences of public databases (IMGT, etc.) and published articles (HTS data) from subjects with physiological or various pathological conditions. Moreover, we analyzed the composition characteristics of the RSSs in human IGHJ genes. Our results suggest that the consensus nonamer and heptamer, the standard spacer length (23 bp), and the mutation site of RSSs may affect the usage frequency of 6 IGHJ gene segments (nonrandom selection), and this specific primary repertoire may result in the lack of significant changes in the usage frequency of 6 IGHJ genes in the peripheral repertoire under physiological and various pathological conditions.

## RESULTS

### The IGHJ gene frequency follows a similar pattern and is rarely influenced by antigen selection

The number of BCR-H sequences from 6 healthy volunteer samples ranged from 250,000 to 1,250,000 (Supplementary Table 1). The order of frequency of IGHJ genes (in-frame) was IGHJ4>IGHJ6>IGHJ3>IGHJ5>IGHJ2>IGHJ1, while out-of-frame sequences followed an order of IGHJ4>IGHJ6>IGHJ5>IGHJ3>IGHJ1>IGHJ2 (Figure 1A). For these two groups, the frequency of IGHJ4 was significantly higher than that of each IGHJ gene, while IGHJ1 and IGHJ2 were significantly less frequently used (Figure 1A). Supplementary Table 2 shows the data of the naive B cell repertoire (primary repertoire, n=48,167) and the memory B cell repertoire (n=50,290). The order of IGHJ gene usage (in-frame) was IGHJ4>IGHJ6>IGHJ3>IGHJ5>IGHJ2>IGHJ1. Sequences (n=9,340) from the IMGT/LIGM-DB also followed this pattern (Supplementary Table 3 and Figure 1C), while the usage of IGHJ genes (out-of-frame) followed IGHJ4>IGHJ6>IGHJ5>IGHJ3>IGHJ1>IGHJ2 (Figure 1B). Similarly, IGHJ4 was significantly used, while the IGHJ1 or IGHJ2 frequency was significantly lower than those of other IGHJ genes.

**Figure 1.**
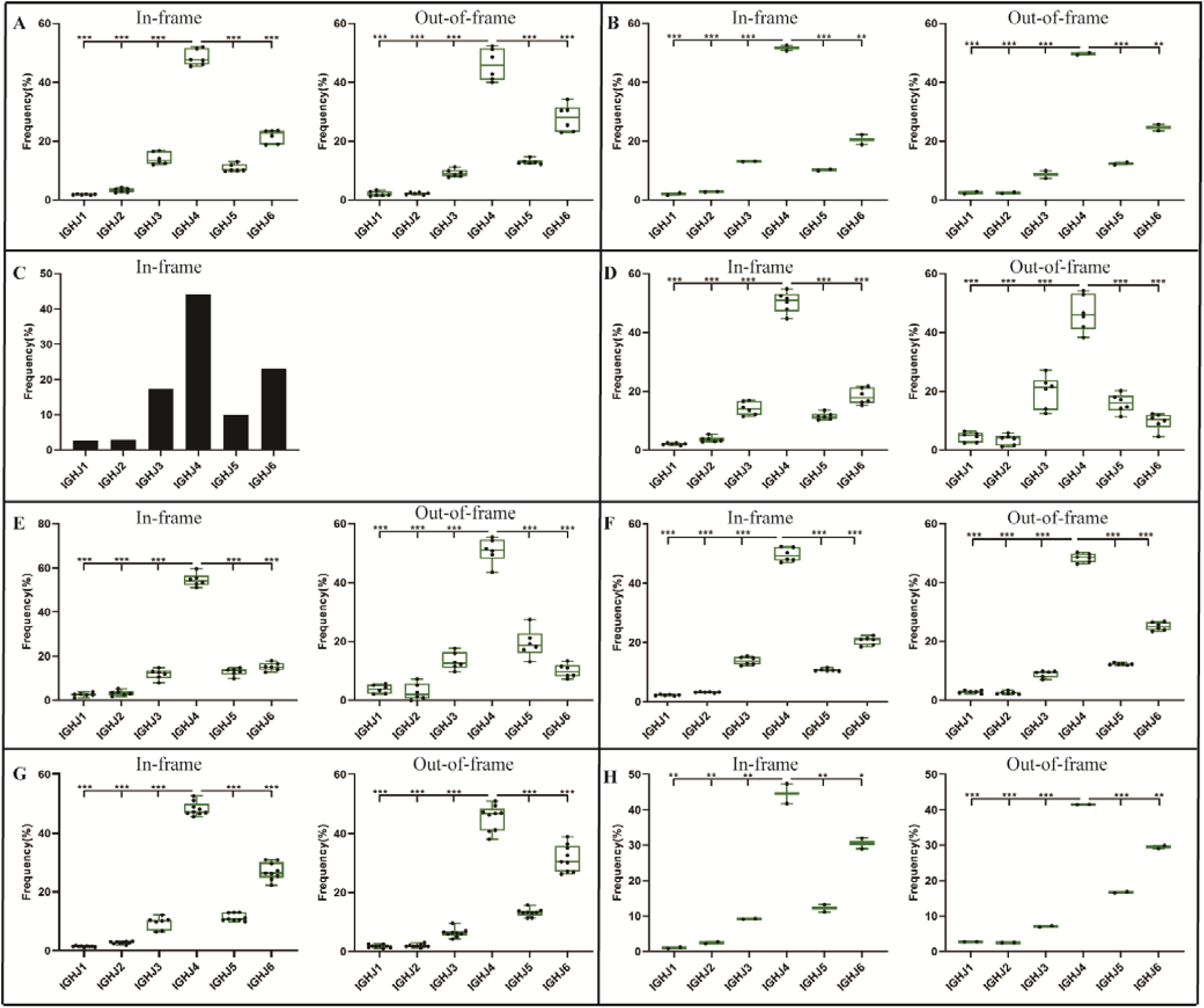
The usage frequencies of 6 IGHJ genes in the in-frame and out-of-frame BCR-H repertoire from different subjects. **(A)** The IGHJ usages of BCR-H repertoire from 6 Healthy volunteers. **(B)** The IGHJ usages of BCR-H repertoire from public data. **(C)** The IGHJ usages of BCR-H repertoire from IMGT data. **(D)** The IGHJ usages of IgM-H repertoire from volunteers before and after immunization with the HBV vaccine. **(E)** The IGHJ usages of IgG-H repertoire from volunteers before and after immunization with the HBV vaccine. **(F)** The IGHJ usages of BCR-H repertoire from SLE volunteers. **(G)** The IGHJ usages of BCR-H repertoire from breast cancer volunteers. **(H)** The IGHJ usages of BCR-H repertoire from volunteers with a high titer of HbsAb.

A similar pattern of IGHJ gene frequency was found not only under physiological conditions but also under pathological conditions. IgM and IgG sequences from three volunteers before and after HBV vaccine are shown in Supplementary Table 4. IgM in-frame sequences presented as IGHJ4>IGHJ6>IGHJ3>IGHJ5>IGHJ2>IGHJ1, while IgM out-of-frame sequences showed IGHJ4>IGHJ3>IGHJ5>IGHJ6>IGHJ1>IGHJ2 (Figure 1D). For IgG sequences, IGHJ4>IGHJ6>IGHJ5>IGHJ3>IGHJ2>IGHJ1 was found in the in-frame sequences, while out-of-frame sequences showed IGHJ4>IGHJ5>IGHJ3>IGHJ6>IGHJ1>IGHJ2 (Figure 1E). The BCR-H sequences from 6 SLE samples ranged from 170,000 to 610,000 sequences (Supplementary Table 5). The usage frequency of 6 IGHJ genes (in-frame) followed IGHJ4>IGHJ6>IGHJ3>IGHJ5>IGHJ2>IGHJ1, while the order of usage frequency of 6 IGHJ genes (out-of-frame) was IGHJ4>IGHJ6>IGHJ5>IGHJ3>IGHJ1>IGHJ2 (Figure 1F). The BCR-H sequence number from breast cancer samples was approximately 70,000∼160,000 for each sample (Supplementary Table 6), and the sequence number from two volunteers with a high titer of HBsAb was 760,000 and 880,000 (Table 7). Interestingly, in-frame and out-of-frame sequences from these two groups consistently presented as IGHJ4>IGHJ6>IGHJ5>IGHJ3>IGHJ2>IGHJ1 (Figure 1G and H).

In addition, we analyzed the ratio of unique to total sequences of each IGHJ gene (in-frame and out-of-frame) and found no differences in 6 IGHJ gene families (Supplementary Table 1 and 2, and Figure 2), which suggests that the multiplex PCR library and the experimental system of HTS did not show obvious bias. Taken together, these results indicate that IGHJ gene frequency follows a similar pattern where IGHJ4 is used at high frequency (>40%), IGHJ6/IGHJ3/IGHJ5 is used at medium frequencies (10%∼20%), and IGH2/IGHJ1 is used at low frequencies (<4%). Therefore, the pattern shows high consistency in physiological and various pathological conditions, which suggests that the recombination selection of each IGHJ gene is nonrandom and rarely influenced by antigen selection.

**Figure 2.**
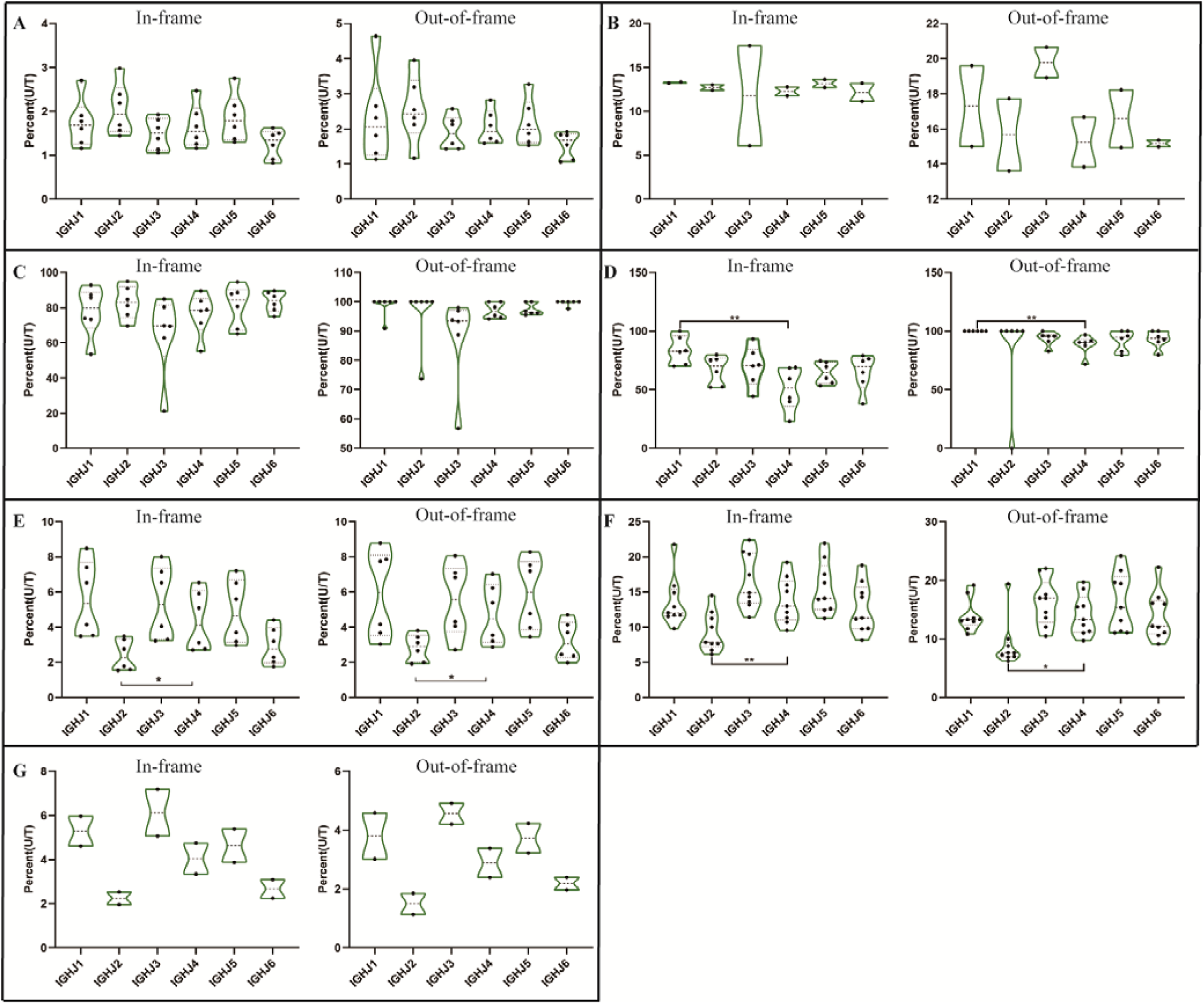
The ratio of unique to total sequences (U/T) of 6 IGHJ genes in the in-frame and out-of-frame BCR-H repertoires from different subjects. **(A)** The IGHJ usages of BCR-H repertoires from 6 Healthy volunteers. **(B)** The IGHJ usages of BCR-H repertoires from public data. **(C)** The IGHJ usages of IgM-H repertoires from volunteers before and after immunization with the HBV vaccine. **(D)** The IGHJ usages of IgG-H repertoires from volunteers before and after immunization with the HBV vaccine. **(E)** The IGHJ usages of BCR-H repertoires from SLE volunteers. **(F)** The IGHJ usages of BCR-H repertoires from breast cancer volunteers. **(G)** The IGHJ usages of BCR-H repertoires from volunteers with a high titer of HbsAb.

### IGHJ-IGHD pairing and trimming and insertion between IGHD and IGHJ

Six IGHJ gene families have different initial BCR-H repertoires, which may be related to nonrandom selection of D-J recombination, thus prompting us to investigate IGHJ-IGHD pairing (Figure 3) and trimming and insertion between IGHD and IGHJ (Figure 4). Most of the 27 IGHD gene subfamilies showed a higher proportion of IGHJ4 pairing (Figure 3). However, whether they were in frame or out of frame, the paired IGHD genes of different IGHJ genes at high or low frequencies were similar. For 6 IGHJ gene families, the IGHD genes paired at high frequency included IGHD6-13, IGHD6-19, IGHD3-22, IGHD3-10, and IGHD2-15, while the low frequency parings included IGHD1-20, IGHD1-7, IGHD4-11, IGHD6-25, and IGHD7-27 (Figure 3).

**Figure 3.**
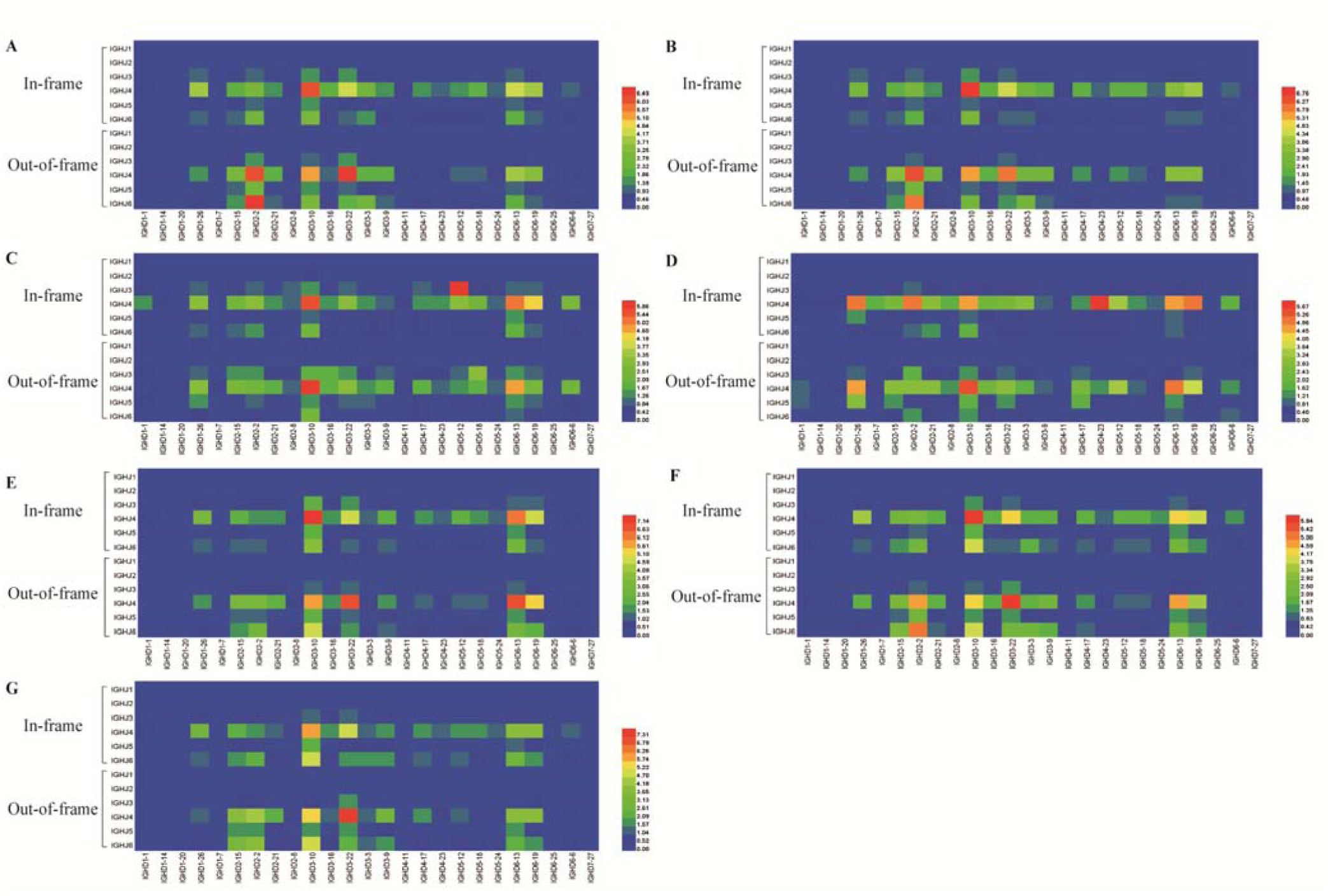
IGHJ-IGHD pairing in the in-frame and out-of-frame BCR-H repertoires from different subjects. **(A)** The IGHJ usages of BCR-H repertoires from 6 Healthy volunteers. **(B)** The IGHJ usages of BCR-H repertoires from public data. **(C)** The IGHJ usages of IgM-H repertoires from volunteers before and after immunization with the HBV vaccine. **(D)** The IGHJ usages of IgG-H repertoires from volunteers before and after immunization with the HBV vaccine. **(E)** The IGHJ usages of BCR-H repertoires from SLE volunteers. **(F)** The IGHJ usages of BCR-H repertoires from breast cancer volunteers. **(G)** The IGHJ usages of BCR-H repertoires from volunteers with a high titer of HbsAb.

**Figure 4.**
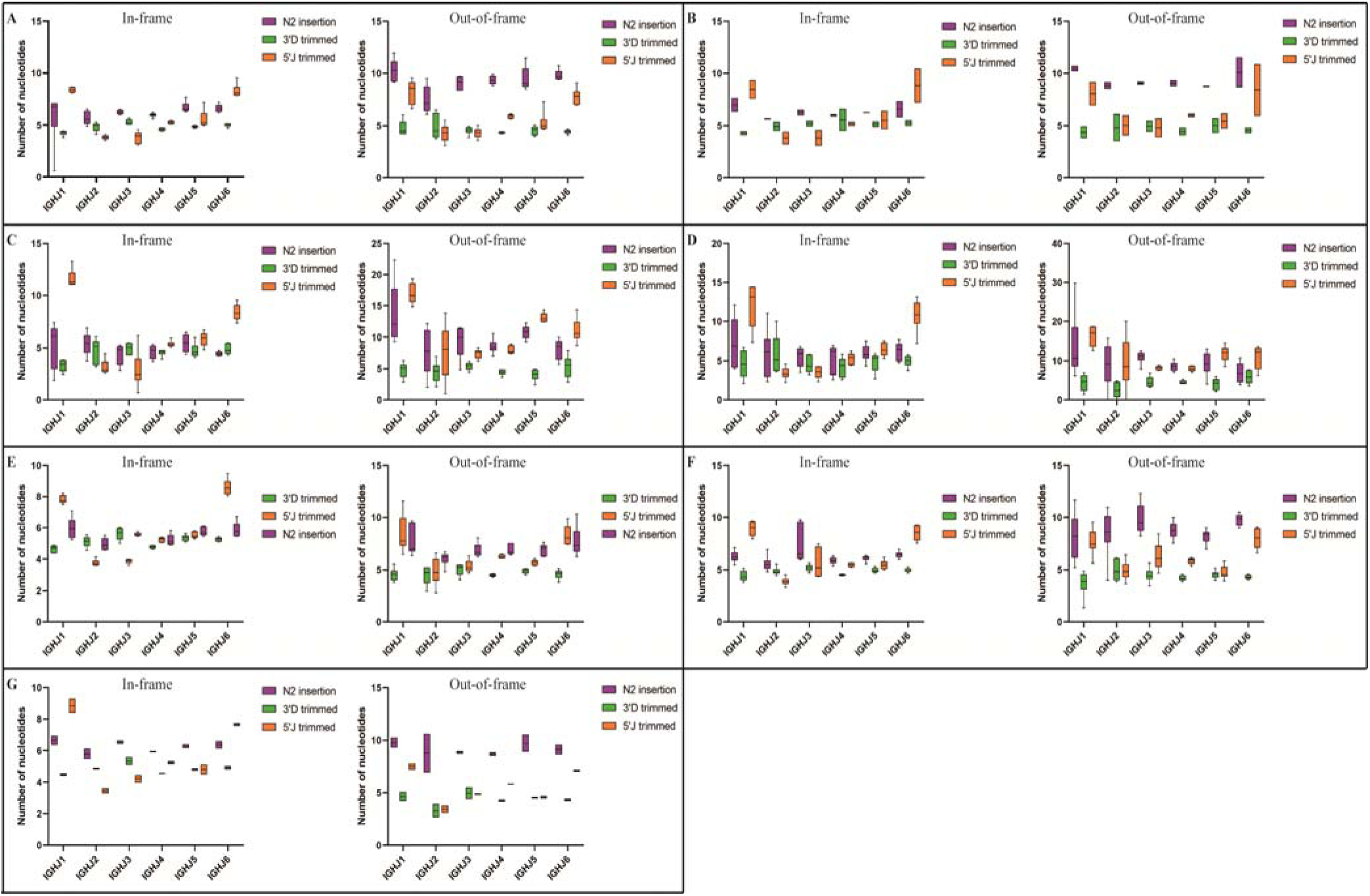
3’D trimmed, 5’J trimmed and N2 insertion at IGHD-IGHJ junction in the in frame and out of frame BCR-H repertoires from different subjects. **(A)** The IGHJ usages of BCR-H repertoires from 6 Healthy volunteers. **(B)** The IGHJ usages of BCR-H repertoires from public data. **(C)** The IGHJ usages of IgM-H repertoires from volunteers before and after immunization with the HBV vaccine. **(D)** The IGHJ usages of IgG-H repertoires from volunteers before and after immunization with the HBV vaccine. **(E)** The IGHJ usages of BCR-H repertoires from SLE volunteers. **(F)** The IGHJ usages of BCR-H repertoires from breast cancer volunteers. **(G)** The IGHJ usages of BCR-H repertoires from volunteers with a high titer of HbsAb.

Trimming and insertion between IGHD and IGHJ mainly presented as 3’D trimmed, 5’J trimmed, and N2 insertion (Figure 4). We found that the mean length of 5’J trimmed showed significant differences among different IGHJ genes under some conditions, while 3’D trimmed and N2 insertion did not show significant differences (data not shown). For IGHJ1 and IGHJ2, the 5’J trimmed length of IGHJ1 (in-frame sequences) showed significant differences compared with the other IGHJ subfamilies in the SLE and IgM with HBV vaccine groups (one-way ANOVA with Bonferroni correction, p<0.001). A similar situation occurred on 5’J trimmed of IGHJ2 in the breast cancer group. The mean length of 5’J trimmed of the IGHJ4 (in-frame or out-of-frame sequences) showed significant differences compared with the other IGHJ genes in the SLE group (one-way ANOVA with Bonferroni correction, each p<0.001). In all groups, IGHJ4 (high usage) showed significant differences compared with IGH1 and IGHJ2 (low usage) (one-way ANOVA with Bonferroni correction, each p<0.001). The mean length of 5’J trimmed from IGHJ6/IGHJ5/IGHJ3 (in-frame sequences) showed significant differences compared with that of the other 5 IGHJ subfamilies in different groups (one-way ANOVA with Bonferroni correction, each p<0.001). These results suggest that the composition of the IGHJ front end (5’J trimmed) may have an impact on the usage and efficiency of the D-J recombination, especially for the IGHJ genes with high or low usage.

### The usage frequency of 6 IGHJ families in the BCR-H repertoires from 19 published articles

We analyzed the usage frequency of the 6 IGHJ gene families in BCR-H repertoires from 19 published articles (29-47) (Supplementary Table 8). Overall, subjects included healthy volunteers of different ages (2 months to 87 years) and patients with different pathological conditions, including SLE, primary biliary cholangitis (PBC), colorectal adenoma and carcinoma (CRC), celiac disease (CD), congenital heart disease, atopic dermatitis, hepatitis C virus infection, rheumatoid arthritis, and primary immune thrombocytopenia, as well as in humanized NOD-scid-IL2R gamma (null) mice. The sample sources included peripheral blood, PBMC (DNA), PBMC (RNA), cord blood, biopsies (RNA), humanized mouse spleen, bone marrow, mucosal tissues, small intestine, lung, stomach, lymph node, tonsil, and thymus. The B cell subsets included B cells, pre-B cells, immature B cells, transitional B cells, naive B cells, normal B cells with IGHV1-69-DJ-C rearrangements, memory B cells, plasmacytes, etc.

The usage frequency of the IGHJ4 gene subfamily was higher than that of other IGHJ genes, suggesting that IGHJ4 had the highest frequency in the initial rearrangement and showed high consistency in peripheral repertoires (after self-tolerance selection or the clonal proliferation response). The usage frequencies of IGHJ1 and IGHJ2 were significantly lower than those of the other IGHJ genes, suggesting that IGHJ1 and IGHJ2 may be partially restricted in the initial rearrangement and that they showed consistency in the peripheral repertoires. IGHJ6, IGHJ3, and IGHJ5 have a medium usage frequency, and the usage frequency of IGHJ6 was higher than that of IGH3 and IGHJ5, except for articles 2, 7, 8, 13 and 19. Additional results showed that IGHJ3 usage was higher than IGHJ5. Regardless of the physiological or pathological conditions, the usage frequencies of the 6 IGHJ gene families in our results are almost identical to those in the 19 published articles. The overall results indicate the nonrandomness of the 6 human IGHJ gene usages during the initial rearrangement process.

### IGHD-IGHJ recombination may affect IGHJ gene usage through the RSS composition

Recombination of IGHJ-IGHD can be divided into two phases. The first phase involves recognition and cleavage of the DNA, and the second phase involves resolution and joining (4, 6). In the evolutionary process, the human IGHJ nonamer sequence is 5’-GGTTTTTTT-3’ (the complementary sequences, CCAAAAACA), and the IGHD nonamer sequence is 5’-ACAAACC-3’ (the complementary sequences, TGTTTTTGG). This evolutionary IGHD-IGHJ “double-stranded complementary pairing” relationship may play a role in the efficiency of D-J recombination. The IGHJ-IGHD recombination schematic diagram is shown in Figure 5A.

**Figure 5.**
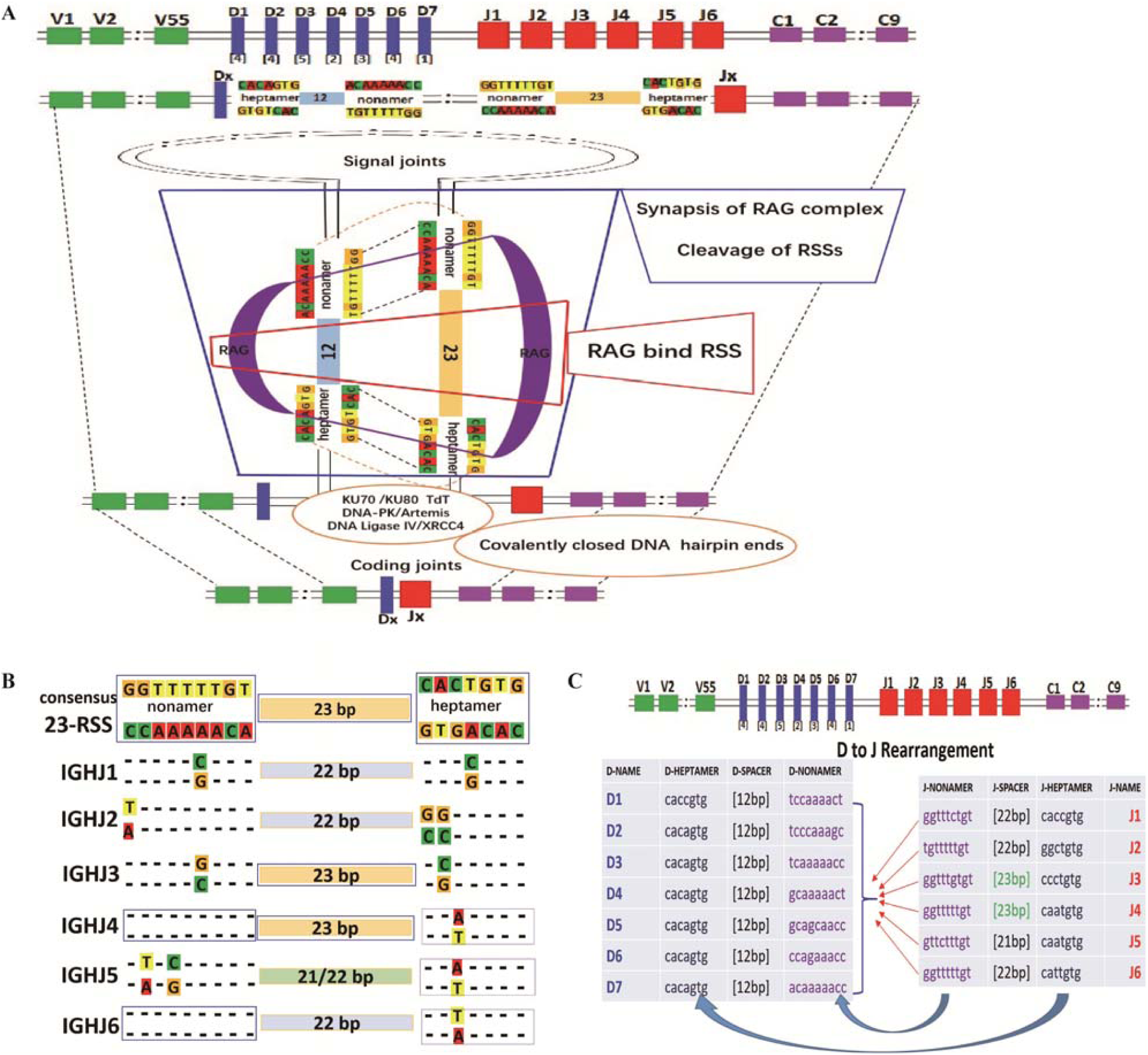
RSS composition characteristics during the IGHD-IGHJ recombination. **(A)** The schematic diagram of IGHJ and IGHD recombination. **(B)** The composition characteristics of human 9-23-7 RSSs (IGHJ-nonamer--IGHJ-spacer--IGHJ-heptamer). **(C)** The pairing of IGHJ (7-12-9) RSSs and IGHD (9-23-7) RSSs during the IGHD-IGHJ recombination.

To investigate whether RSSs affect IGHJ usage, we obtained human IGHJ gene sequences (X97051, X86356, M25625, J00256, AJ879487 from the IMGT and GenBank) for RSS composition analysis. The composition and characteristics of the human IGHJ RSSs (nonamer--spacer--heptamer (9-23-7)), J region sequence and AA are shown in Supplementary Tables 9-11. IGHJ4 and IGHJ6 have the consensus nonamer sequences “5’-GGTTTTTGT-3’” (the complementary sequence is “CCAAAAACA”). However, the nonamer had one or two base mutations in other IGHJ families. Position 4 of IGHJ1 mutated from A to G, position 9 of IGHJ2 mutated from C to A, position 4 of IGHJ3 mutated from A to C, position 6 of IGHJ5 mutated from A to G, and position 8 of IGHJ5 mutated from C to A (Supplementary Table 9 and Figure 5B). The consensus heptamer is CACAGTG/GTGTCAC. Position 5 of IGHJ4 and IGHJ5 mutated from G to T (IGHJ6 mutated to A), position 4 of IGHJ1 mutated from A to G, position 6 of IGHJ2 mutated from T&G to C, and position 6 of IGHJ3 mutated from T to G. In addition, IGHJ4 and IGHJ3 have a consensus spacer length (23 bp), while the spacer length is reduced by 1 or 2 bases in other IGHJ gene families (IGHJ1-22 bp, IGHJ2-22 bp, IGHJ5-21 or 22 bp, and IGHJ6-22 bp) (Figure 5C).

Overall, compared to the conserved RSS, the IGHJ4 gene subfamily is roughly consistent, the spacer lengths are changed in IGHJ6, the nonamer and heptamer are altered in IGHJ3, the spacer lengths and the nonamer are changed in IGHJ5, and the nonamer, heptamer, and spacer lengths are changed simultaneously in IGHJ1 and IGHJ2. There were different code end sequences (AA) in the IGHJ genes IGHJ4 (15AA), IGHJ1 and IGHJ2 (17AA), IGHJ3 and IGHJ5 (16AA), and IGHJ6 (20AA).

## DISCUSSION

The V(D)J gene family of the human BCR heavy chain variable region contains 56 functional V genes with 3’ ends of 7-23-9 RSS, 27 functional D genes with 3’ ends of 9-12-7 RSS and 5’ ends of 7-12-9 RSS, and 6 functional J genes with 5’ ends of 9-23-7 RSS. The recombination starts with recombination of the 3’ end of the D gene and the 5’ end of the J gene, and then the 3’ end of the V gene is recombined with the 5’ end of the D gene (D-J recombination). In the peripheral BCR-H repertoires, the usage frequency of each V(D)J gene is related to the preferred usage in the initial rearrangement, the selection of self-tolerance and the response of peripheral clonal proliferation. However, the mechanism and significance of differential selection among V(D)J gene subfamilies have not been fully elucidated (4, 6, 48).

We investigated the usage frequency of the 6 IGHJ genes in unique BCR-H repertoires (in-frame and out-of-frame) by HTS under physiological and various pathological conditions. In addition, we analyzed non-HTS-derived BCR-H sequences from the IMGT database, the HTS-derived BCR-H sequences from the public database (other laboratory), and the usage frequency data of 6 IGHJ genes from 19 published articles. The results indicate that IGHJ4 has a significantly high usage frequency in all subjects, various tissues, and different B cell subset samples. IGHJ6, IGHJ3, and IGHJ5 have medium usage frequencies, and IGHJ1 and IGHJ2 have significantly low usage frequencies. Taken together, these results suggest that the recombination selection of each human IGHJ gene is nonrandom and rarely influenced by antigen selection, which is different from the traditional understanding.

### The IGHJ nonamer and combination frequency

Early studies suggested that the composition characteristics of human IGHJ RSSs may affect the usage frequency of IGHJ in the initial rearrangement. In 1987, Akira S et al found that two sets of heptamer (CACTGTG) and nonamer (GGTTTTTGT) sequences were enough to initiate the V(D)J joining if the 12-bp and 23-bp spacer rule is satisfied in the recombination-competent pre-B cell line (49). A point mutation in the heptamer sequence or a change in the combination of the two spacer lengths (21 bp 22 bp24 bp/11 bp13 bp) would drastically reduce the recombination frequency.

Variation from the conserved sequences in the heptamer and nonamer of the RSSs is considered a major factor affecting the relative representation of gene segments in the primary repertoire. The mechanism of RSSs on gene recombination is mainly related to the interaction efficiency of RAG protein (recombinase) (50-54). Based on the composition of human IGHJ gene families, we found differences in RSSs among 6 IGHJ gene families (Supplementary Table 9 and Figure 5), which suggests that these differences may affect the usage frequency (nonrandom) of IGHJ gene families.

The nonamer of human IGHJ4 and IGHJ6 is the conserved sequence 5’-GGTTTTTGT-3’ or 5’-CCAAAACA-3’, while the other IGHJ nonamers have one or two base mutations. Experiments in vitro based on B cell lines showed that the mutation of nonamer had a significant effect on the corresponding gene recombination. Ramsden DA et al found that the nonamers were probably the most important element in initial RAG protein binding (12). A single base mutation of the nonamer resulted in a reduction in overall cleavage levels when the heptamer was retained, but the entire nonamer was substituted with random sequence. Both nicks and hairpins were still found, but overall cleavage was reduced fold. Kowalski D et al found (55, 56) that A-rich core sequences of the nonamer may be important to facilitate strand dissociation during the process of recombination.

The presence of three consecutive A residues was necessary for efficient recombination in the nonamer; furthermore, the nucleotides flanking the A-rich core needed to be other than one residue. The mechanism may be that the recombinase must measure the distance between the heptamer and the nonamer to satisfy the 12/23-bp spacer rule (3, 12, 13, 14, 15). Akamatsu Y et al found that the A residue at position 5 (nonamer A-rich core) was most crucial in their recombination assay (13). However, Hesse JE et al considered that the “A residue” at position 6 (nonamer) was most crucial in their recombination assay (3). Regarding the effect of nonamer A-rich core mutation and corresponding gene usage, Akamatsu Y et al found that recombination frequency decreased to 27.3% of the control with the mutant 9-4G (position 4 was changed to G) (13). A mutant at position 9-5C gave the lowest recombination frequency (10.4%). With the double mutant at positions 9-3G and 9-4G, the joining rate dropped only to 19.3% (9-6G and 9-7G was 26.0%). According to the results from cell line experiments, human IGH4 and IGHJ6 gene subfamilies appear to have a “complete A-rich core” in the nonamer (conserved), which may play an important role in their high usage selection. However, 9-4A of human IGHJ1 is mutated to 9-4G, 9-4A of IGHJ3 is mutated to 9-4C, 9-6A of IGHJ5 is mutated to 9-6G, and 9-8C is mutated to 9-8A, which is a possible cause of their disfavored usages.

In addition, Akamatsu Y et al found that the nonamer 9-2C was changed to 9-2A, and the recombination frequency was reduced to 2.7% of the control level; 9-2C was changed to 9-2T, and the frequency was reduced to 12.9%; and 9-2C was changed to 9-2G, and the frequency remained at 61.3% (13). When 9-8C/9-9C were changed to 9-8N/9-9A, the recombination frequency dramatically dropped to less than 0.1%, which suggested that the C residue plays an important role when the recombinase measures the distance between the heptamer and the nonamer sequences. In this study, one factor for the low usage frequency of the human IGHJ2 gene may be its 9-9C mutation to 9-9A.

### The IGHJ heptamer and combination frequency

Human IGHJ4 and IGHJ5 genes have the same heptamer sequence (CAATGTG/GTTACAC). Position 7-3C is mutated to 7-3A compared to the conserved heptamer, and 7-3C is mutated to 7-3T in IGHJ6, while the heptamer sequences of the IGHJ4/IGHJ5/IGHJ6 gene subfamilies are uniform on the double strand. Position 7-4A is mutated to 7-4G in the IGHJ1 gene, position 7-6T/7-7G is mutated to 7-6C/7-7C in the IGHJ2 gene, and position 7-6T is mutated to 7-6G in the IGHJ3 gene.

The relationship between the heptamer and the recombination frequency of the corresponding gene family has been confirmed by several laboratories. Both studies found that the mutation of the entire heptamer resulted in low levels of nicking distributed across several sites, the mechanism of heptamer affecting recombination was related to the formation of hairpins, and the nicks and hairpins were reduced 2-fold when the sequence of the last four positions of the heptamer was changed (12, 57). In addition, nicking formation depended on the heptamer for the generation of double strand breaks (DSBs) by RAG1 and RAG2, and the nonamer at the correct distance would improve heptamer efficiency in the natural RSSs. The first three nucleotide positions were nearly 100% conserved (CAC/GTG) in the BCR gene. The mutations were in the first three positions, and cleavage was impaired either at the nicking step or the hairpin formation site. No rearrangement was detected with the mutant at position l (7-1G). Mutations at position 2 (7-2T) and position 3 (7-3G) produced detectable levels of recombination, 0.5% and 0.6%, respectively. The G residue at position 5 was changed to C (7-5C), and the recombination frequency dropped to 5.9% of the control level. For the rest of the residues in the heptamer, mutation effects were moderate, ranging from 28.5 to 52.0%. Akamatsu Y et al found that no rearrangement was detected with the mutant at position l (7-1G), and mutations at position 2 (7-2T) and position 3 (7-3G) produced detectable levels of recombination, 0.5% and 0.6%, respectively. The recombination frequency dropped to 5.9% of the control level when the G residue at position 5 was changed to C (7-5C); for the rest of the residues in the heptamer, mutation effects were moderate, ranging from 28.5 to 52.0% (13).

The first three positions of the 6 human IGHJ gene subfamily heptamers are a conserved CAC/GTG sequence. Based on the results of Akamatsu Y et al, position 7-4A of human IGHJ1 mutated to 7-4G, and 7-6T/7-7G of IGHJ2 mutated to 7-6C/7-7C, which may be one important factor causing their low usage frequency. In addition, the 7-5G mutation (IGHJ3, IGHJ4, IGHJ5, and IGHJ6) may have a moderate effect on their usage frequency. The effect of mutations in the human IGHJ heptamer on usage frequency needs to be further explored.

### The RSS spacer and combination frequency

The length of the spacer is also a determining factor contributing to the usage frequency of V(D)J rearrangement. Human IGHJ4 and IGHJ3 gene subfamilies have a conserved 23 bp length; however, the IGHJ1, IGHJ2, IGHJ5, and IGH6 gene subfamilies have 21 bp or 22 bp spacer lengths. Akamatsu Y et al found that the recombination frequency dropped to 7.7% with the 11-bp RSSs when one C residue was added to the 12 bp RSSs (13 bp spacer) (11.0% joining rate); when two C residues were added (14 bp spacer), recombination dropped below the detection level, indicating that RSS spacer length was critical for combination frequency (13). Nadel B et al found that the effect of the spacer on the recombination rate of various human Vk gene segments in the peripheral repertoire correlated with their frequency in pre-B cells (in vivo) (16). Steen SB et al found that changing the spacer length by one nucleotide (23 bp1 bp only moderately reduced DSB formation, altering the spacer length by greater than one nucleotide (23 bp-2 bp and 23 bp-3 bp), severely reduced cleavage to a lesser degree (15). If each RSS contains a severe mutation (12 bp-3 bp/23 bp-3 bp), no DSBs were observed. According to the above research, the length of the 23 bp spacer of the human IGHJ4 and IGHJ3 gene subfamilies is an important factor in the higher usage frequency, and the length reduction of the 23 bp spacer in the IGHJ1, IGHJ2, IGHJ5, and IGHJ6 genes reduces their recombination usage.

The sequences of RSSs may affect the usage frequency of V(D)J gene recombination. Fanning L et al found that when the Igk 12 bp spacer of the natural sequence CTAC “A” GACTGGA was changed to CTAC “C” GACTGGA but the corresponding 23RSSs-GTAGTACTCCACTG TCTGGCTGT were not changed, the mutant proximal RSSs were consistently used less frequently (17). In addition, the recombination efficiency was 63.0% of the control level when the 12 bp spacer was changed to an artificial sequence GATCGATCGATC (13, 57, 58). Larijani M et al found that the frequency of recombination decreased by approximately 5-fold when the V81x spacer (AGCAAAAGTTACTGTGAGCTCAA) was replaced by that of VA1 (TTGTAA CCACATCCTGAGTGTGT) (14). Montalbano A et al found that single base pair changes in the spacer sequence can significantly affect recombination efficiency (18). Nadel B et al confirmed that natural variation in spacer sequences could contribute to the nonrandom use of human V genes observed in vivo and that a randomly generated variant of a human V spacer was significantly worse in recombination efficiency (16). These results suggest that the spacer sequence plays an important role in recombination efficiency. Our results show that the ratio of AT and CG in 23 bp spacer sequences of 6 human IGHJ gene families is inconsistent (Supplementary Table 9). Base C has the highest ratio in IGHJ4. Is this the reason for the higher usage frequency in the recombination of the IGHJ4 gene subfamily? Whether the base composition of spacer sequences such as the nonamer has the key “A-rich core” structure need to be further explored.

### Distance and combination frequency

It has been confirmed that the proximal gene has preferred usage in the initial rearrangement (8-10). Malynn, B.A., et al. believe that the difference in IGHV gene usage in adult spleen B cells is mainly due to the selection of the initial rearrangement rather than the changes in expression frequency after rearrangement (59). The “proximal and distal” studies of BCR recombinant genes are mainly focused on the V gene. “Proximal and distal” differences in the J gene have not been reported. In our results, we did not find the “proximal” phenomenon in the 6 IGHJ gene families with high usage frequencies.

### Other factors and combination frequency

Ramsden DA et al found that the sequence of the coding end may be related to the usage frequency of gene combination (12). We found that there are differences in the amino acid length and the coding flank sequences of the human 6 IGHJ families (Supplementary Table 9). The IGHJ4 gene has the shortest 16 amino acid components. The sequence of the coding end and AA length may affect the usage frequency of IGHJ. We analyzed the deletions of the 3’D end, the 5’J end and the insertion between the D-J end and found that there was a difference between the 5’J end of IGHJ4 and other IGHJ genes (Figure 3 and Supplementary Table 11). Whether it was a factor for high usage of IGHJ4 needs to be further studied. In addition, IGHD gene families may also affect the nonrandomness of IGHJ genes. VanDyk LF et al suggested that V(D)J recombination was targeted by RSSs, while the RSSs flanking D segments appeared to be equivalent. They were not randomly utilized, suggesting that the D-3’ RSSs were not simply superior targets for the D-J recombinase but instead that targeting certain 12/23-bp spacer RSS combinations is more effective (60).

We found that the conserved nonamer of IGHJ4 and IGHJ6 had a higher “double-stranded complementary paired” rate than the 27 IGHD nonamer sequences (Supplementary Table 10 and Supplementary figure 1), although it did not show obvious differences. At present, no evidence to support that RAG has an effect on the “double-stranded complementary paired” of the J-heptamer to D-heptamer and J-nonamer to D-nonamer exists; the mechanism is still unknown. We hypothesize that two genes with high complementarity (7-7/9-9) may be more favorable for binding, cleaving, hairpin formation, and DSB in the recombination process (Figure 5), which is a very interesting entry point for further research in BCR gene recombination.

In summary, for the possible impacts of RSSs on IGHJ usage frequency, RSSs of human IGHJ4 genes are consistent with conserved RSSs. The length of the IGHJ6 spacer (23 bp) changed slightly, the nonamer and heptamer of IGHJ3 changed, and the length and nonamer of IGHJ5 changed. However, the nonamer, heptamer and spacer of IGHJ1 and IGHJ2 changed significantly. These may be factors that resulted in nonrandom usage of the human IGHJ gene (generally, IGHJ4>IGHJ6>IGHJ3> or ≈IGHJ5>IGHJ2≈IGHJ1) in the initial rearrangement. In the initial human BCR-H repertoires (before antigen selection), the “background” of the 6 IGHJ genes (the initial usage frequency) is different and rarely influenced by antigen selection. These results suggest that re-evaluation and further investigation are needed when analyzing the significance and mechanism of each IGHJ gene usage in self-tolerance selection and the clonal proliferative response.

## MATERIALS AND METHODS

### Subjects and sample preparation

The subjects included six healthy volunteers (6 samples: H-1, H-2, H-3, H-4, H-5 and H-6) (22), two volunteers with systemic lupus erythematosus (SLE) (including 6 total samples pretreatment, during treatment and after treatment, namely, S1-1, S1-2, S1-3, S2-1, S2-2 and S2-3) (23), three volunteers with breast cancer (including 9 total samples pretreatment, during treatment and after treatment, namely, B3-1, B2-1, B1-1, B3-2, B2-2, B1-2, B3-3, B2-3, and B1-3), two volunteers with a high titer of HBsAb (2 samples: HBsAb-1, HBsAb-2) (24) and three volunteers with samples before and after immunization with the HBV vaccine (6 IgM samples (V1-BM, V1-AM, V2-BM, V2-AM, V3-BM, V3-AM) and 6 IgG samples (V1-BG, V1-AG, V2-BG, V2-AG, V3-BG, and V3-AG)) (25). The peripheral blood samples were obtained from the Affiliated Hospital of Zunyi Medical University. All the volunteers were informed of the purpose of peripheral blood collection and were under a protocol approved by The Committee on the Ethics of Human Experiments of Zunyi Medical University, and all the experiments were performed in accordance with the guidelines of the committee. Peripheral blood mononuclear cells (PBMCs) were obtained using Ficoll 1640 (Biochrom AG, Berlin, Germany) density centrifugation.

### Total RNA/DNA extraction and cDNA synthesis

Total RNA was extracted from the PBMCs in three volunteers with immunization with HBV vaccine according to the manufacturer’s protocol for the total RNA extraction kit (OmegaBio-Tek). The total RNA was then reverse transcribed into cDNA using Oligo dT18 according to the manufacturer’s protocol for the reverse transcription kit (MBI, Fermentas). The genomic DNA from PBMCs in other samples was obtained using the QIAamp DNA Mini Kit (QIAGEN, CA) and was stored in a QIAsafe DNA tube (QIAGEN).

### High-throughput sequencing

All the DNA samples were sent to Adaptive Biotechnologies Corp (http://www.adaptivebiotech.com) for multiplex PCR amplification of human BCR-HCDR3 regions. Error from bias in this multiplex PCR assay was controlled using synthetic templates (26), and the HCDR3 sequences were acquired by HTS on the ImmunoSEQ platform (http://www.adaptivebiotech.com) (23). All the PCR products of cDNA samples after PCR amplification were sent to Tongji-SCBIT Biotechnology Corporation for HTS, and detailed experimental procedures have been described in our previous article (25). The HCDR3 regions were identified within the sequencing reads according to the definition established by the International ImMunoGeneTics (IMGT) collaboration. A standard algorithm was used to identify which V(D)J segments contributed to each HCDR3 sequence.

### Public data

We used 9,340 unique in-frame BCR-H sequences (non-HTS data in different pathological states) derived from the IMGT/LIGM-DB to analyze the IGHJ gene frequency by IMGT/HighV-QUEST (27). In addition, 50,290 BCR-H sequences of memory B cells and 48,167 HCDR3 sequences of naive B cells from a public database (HTS data from 3 healthy volunteers) were used for this study (28). The unique in-frame BCR-H sequences (n=84,804) and out-of-frame sequences (n=13,653) were compared and analyzed by IMGT/HighV-QUEST software in this study.

### Sequence analysis

The raw sequences in FASTA format were analyzed with IMGT/HighV-QUEST online software (version 1.3.1, http://www.imgt.org). Using the IMGT summary document, the sequences not meeting the following criteria were filtered out: (1) no results (sequences for which IMGT/HighV-QUEST did not return any result) and (2) unknown (sequences for which no functionality was detected. This category corresponds to the sequences for which the junction could not be identified (no evidence of rearrangement, no evidence of junction anchors).). In-frame and out-of-frame unique sequences remaining after filtering were used for IGHJ gene frequency, D-J pairing, and nucleotide insertion and deletion analyses.

### RSS composition analysis

According to the accession numbers of these human IGHJ and IGHD genes in IMGT and GenBank, we obtained detailed annotations of complete human genome sequences for RSS composition analysis, including sequence characteristics of nonamers and heptamers, length characteristics of 12 bp and 23 bp spacers, and the IGHJ gene segment (amino acid) composition of code end.

### Software and statistics

IMGT/HighV-QUEST (version 1.3.1) was used for identification of sequences (JH and DH), evaluation of functionality and statistical analysis of the sequence data; IMGT/V-QUEST (version 3.3.1) was used for identification of nonamers, heptamers, 12 bp and 23 bp spacers, and IGHJ gene segments of the coding end; Microsoft Office Excel (version 365) was used for storage, filtering and statistical analysis of the sequences. The resulting sequences were graphed using Prism 8 software (GraphPad). IGHJ gene usages were compared using a χ2 test. Insertions and deletions of the nucleotides were compared using one-way ANOVA with Bonferroni correction. All statistically significant differences are indicated as *=p<0.05; **=p<0.01, and ***=p<0.001.

## DATA AVAILABILITY

Our sequencing data was deposited in ImmnunoSEQ database (https://clients.adaptivebiotech.com/login). Sequence for RSS analysis included X97051, X86356, M25625, J00256, AJ879487 from the IMGT/LIGM-DB and GenBank.

## ACKNOWLEDGEMENTS

We are grateful to all the volunteers for supporting this study, and we thank all the authors, IMGT and GenBank for using their HTS data for analysis.

