## Supplementary Tables for "The usage of human IGHJ genes follows a particular nonrandom selection: The recombination signal sequence affects the usage of human IGHJ genes"

**Supplementary Table 1.** The sequence data of 6 IGHJ gene families in the in-frame and out-of-frame BCR-H repertoire from 6 Healthy volunteers

| **Sample** | **Repertoire** | | | **In-frame** | | | **IGHJ1** | | | **IGHJ2** | | | **IGHJ3** | | | **IGHJ4** | | | **IGHJ5** | | | **IGHJ6** | | |
| --- | --- | --- | --- | --- | --- | --- | --- | --- | --- | --- | --- | --- | --- | --- | --- | --- | --- | --- | --- | --- | --- | --- | --- | --- |
|  | **total** | | **unique** | **total** | **unique** | | **total** | | **unique** | **total** | **unique** | | **total** | | **unique** | **total** | | **unique** | **total** | | **unique** | **total** | | **unique** |
| H-1 | 550082 | | 6351 | 474996 | 5266 | | 6977 | | 90 | 12735 | 184 | | 59981 | | 634 | 194010 | | 2432 | 49685 | | 681 | 151608 | | 1245 |
| H-2 | 556813 | | 12353 | 482531 | 10455 | | 7485 | | 202 | 14409 | 429 | | 70220 | | 1352 | 219959 | | 5438 | 38288 | | 1053 | 132170 | | 1981 |
| H-3 | 765659 | | 14690 | 658273 | 12285 | | 11000 | | 209 | 14764 | 353 | | 112124 | | 2028 | 300258 | | 5835 | 55841 | | 1190 | 164286 | | 2670 |
| H-4 | 1227335 | | 20722 | 1052967 | 17223 | | 15791 | | 280 | 20565 | 449 | | 177833 | | 2894 | 471857 | | 7832 | 89786 | | 1725 | 277135 | | 4043 |
| H-5 | 499983 | | 5777 | 436701 | 4889 | | 8296 | | 96 | 11744 | 184 | | 60807 | | 688 | 217167 | | 2515 | 37455 | | 487 | 101232 | | 919 |
| H-6 | 897745 | | 12839 | 763705 | 10621 | | 12348 | | 198 | 15385 | 260 | | 96228 | | 1326 | 360141 | | 5072 | 76626 | | 1265 | 202977 | | 2500 |
| Total | 4497617 | | 72732 | 3869173 | 60739 | | 61897 | | 1075 | 89602 | 1859 | | 577193 | | 8922 | 1763392 | | 29124 | 347681 | | 6401 | 1029408 | | 13358 |
| U/T | 72732/4497617 | | | 60739/3869173 | | | 1075/61897 | | | 1859/89602 | | | 8922/577193 | | | 29124/1763392 | | | 6401/347681 | | | 13358/1029408 | | |
| **Sample** | **Repertoire** | | | **Out-of-frame** | | | **IGHJ1** | | | **IGHJ2** | | | **IGHJ3** | | | **IGHJ4** | | | **IGHJ5** | | | **IGHJ6** | | |
|  | **total** | **unique** | | **total** | | **unique** | **total** | **unique** | | **total** | | **unique** | **total** | **unique** | | **total** | **unique** | | **total** | **unique** | | **total** | **unique** | |
| H-1 | 550082 | 6351 | | 75086 | | 1085 | 718 | 13 | | 1269 | | 27 | 5865 | 84 | | 32189 | 527 | | 10311 | 158 | | 24734 | 276 | |
| H-2 | 556813 | 12353 | | 74282 | | 1898 | 668 | 31 | | 1189 | | 47 | 6851 | 176 | | 34443 | 972 | | 7056 | 231 | | 24075 | 441 | |
| H-3 | 765659 | 14690 | | 107386 | | 2405 | 2493 | 66 | | 1254 | | 40 | 12011 | 268 | | 41385 | 991 | | 11932 | 308 | | 38311 | 732 | |
| H-4 | 1227335 | 20722 | | 174368 | | 3499 | 2551 | 59 | | 2473 | | 63 | 16117 | 344 | | 66495 | 1397 | | 20752 | 439 | | 65980 | 1197 | |
| H-5 | 499983 | 5777 | | 63282 | | 888 | 1145 | 15 | | 1631 | | 19 | 5253 | 75 | | 29222 | 465 | | 6806 | 111 | | 19225 | 203 | |
| H-6 | 897745 | 12839 | | 134040 | | 2218 | 6352 | 72 | | 2116 | | 49 | 11306 | 179 | | 54900 | 949 | | 15562 | 291 | | 43804 | 678 | |
| Total | 4497617 | 72732 | | 628444 | | 11993 | 13927 | 256 | | 9932 | | 245 | 57403 | 1126 | | 258634 | 5301 | | 72419 | 1538 | | 216129 | 3527 | |
| U/T | 72732/4497617 | | | 11993/628444 | | | 256/13927 | | | 245/9932 | | | 1126/57403 | | | 5301/258634 | | | 1538/72419 | | | 3527/216129 | | |

Note: U/T represents the ratio of unique to total sequences.

**Supplementary Table 2.** The sequence data of 6 IGHJ gene families in the in-frame and out-of-frame BCR-H repertoire from public data (Naive and Memory B cells)

|  | **Repertoire** | | **In-frame** | | **IGHJ1** | | **IGHJ2** | | **IGHJ3** | | **IGHJ4** | | **IGHJ5** | | **IGHJ6** | |
| --- | --- | --- | --- | --- | --- | --- | --- | --- | --- | --- | --- | --- | --- | --- | --- | --- |
| **Sample** | **total** | **unique** | **total** | **unique** | **total** | **unique** | **total** | **unique** | **total** | **unique** | **total** | **unique** | **total** | **unique** | **total** | **unique** |
| PLOS-1 | 354494 | 48167 | 301527 | 40455 | 4797 | 640 | 8684 | 1077 | 30191 | 5271 | 160720 | 20514 | 29304 | 3997 | 67831 | 8956 |
| PLOS-2 | 398162 | 50290 | 363046 | 44349 | 7939 | 1049 | 9395 | 1224 | 96345 | 5835 | 198216 | 23315 | 36045 | 4583 | 75106 | 8343 |
| Total | 752656 | 98457 | 664573 | 84804 | 12736 | 1689 | 18079 | 2301 | 126536 | 11106 | 358936 | 43829 | 65349 | 8580 | 142937 | 17299 |
| U/T | 98457/752656 | | 84804/664573 | | 1689/12736 | | 2301/18079 | | 11106/126536 | | 43829/358936 | | 8580/65349 | | 17299/142937 | |
|  | **Repertoire** | | **Out-of-frame** | | **IGHJ1** | | **IGHJ2** | | **IGHJ3** | | **IGHJ4** | | **IGHJ5** | | **IGHJ6** | |
| **Sample** | **total** | **unique** | **total** | **unique** | **total** | **unique** | **total** | **unique** | **total** | **unique** | **total** | **unique** | **total** | **unique** | **total** | **unique** |
| PLOS-1 | 354494 | 48167 | 52967 | 7712 | 1034 | 155 | 1220 | 166 | 3003 | 568 | 27883 | 3857 | 6604 | 986 | 13223 | 1980 |
| PLOS-2 | 398162 | 50290 | 35116 | 5941 | 837 | 164 | 852 | 151 | 2833 | 585 | 17573 | 2929 | 3907 | 712 | 9114 | 1400 |
| Total | 752656 | 98457 | 88083 | 13653 | 1871 | 319 | 2072 | 317 | 5836 | 1153 | 45456 | 6786 | 10511 | 1698 | 22337 | 3380 |
| U/T | 98457/752656 | | 13653/88083 | | 319/1871 | | 317/2072 | | 1153/5836 | | 6786/45456 | | 1698/10511 | | 3380/22337 | |

Note: U/T represents the ratio of unique to total sequences.

**Supplementary Table 3.** The sequence data of 6 IGHJ gene families in the in frame and out of frame BCR-H repertoire from IMGT data

| **Unique sequence** | **IGHJ1** | **IGHJ2** | **IGHJ3** | **IGHJ4** | **IGHJ5** | **IGHJ6** |
| --- | --- | --- | --- | --- | --- | --- |
| 9340 | 245 | 270 | 1607 | 4118 | 942 | 2158 |

Note: U/T represents the ratio of unique to total sequences.

**Supplementary Table 4.** The sequence data of 6 IGHJ gene families in the in-frame and out-of-frame BCR-H repertoire (IgM & IgG) from volunteers before and after immunization with the HBV vaccine

| **IgM** | **Repertoire** | | **In-frame** | | **IGHJ1** | | **IGHJ2** | | **IGHJ3** | | **IGHJ4** | | **IGHJ5** | | **IGHJ6** | | **Out-of-frame** | | **IGHJ1** | | **IGHJ2** | | **IGHJ3** | | **IGHJ4** | | **IGHJ5** | | **IGHJ6** | |
| --- | --- | --- | --- | --- | --- | --- | --- | --- | --- | --- | --- | --- | --- | --- | --- | --- | --- | --- | --- | --- | --- | --- | --- | --- | --- | --- | --- | --- | --- | --- |
| **Sample** | **total** | **unique** | **total** | **unique** | **total** | **unique** | **total** | **unique** | **total** | **unique** | **total** | **unique** | **total** | **unique** | **total** | **unique** | **total** | **unique** | **total** | **unique** | **total** | **unique** | **total** | **unique** | **total** | **unique** | **total** | **unique** | **total** | **unique** |
| V1-QM | 689 | 569 | 564 | 447 | 8 | 7 | 16 | 13 | 73 | 51 | 290 | 229 | 58 | 51 | 119 | 94 | 125 | 123 | 3 | 3 | 2 | 2 | 30 | 28 | 65 | 65 | 14 | 14 | 11 | 11 |
| V1-HM | 2275 | 1853 | 1904 | 1497 | 56 | 30 | 54 | 41 | 225 | 181 | 1048 | 821 | 223 | 180 | 298 | 244 | 371 | 356 | 22 | 20 | 19 | 14 | 51 | 50 | 169 | 166 | 67 | 64 | 43 | 42 |
| V2-QM | 994 | 678 | 834 | 524 | 15 | 11 | 22 | 20 | 113 | 71 | 478 | 264 | 109 | 71 | 97 | 87 | 160 | 154 | 7 | 7 | 7 | 7 | 33 | 32 | 74 | 70 | 32 | 31 | 7 | 7 |
| V2-HM | 684 | 619 | 563 | 499 | 14 | 12 | 20 | 19 | 99 | 84 | 249 | 223 | 56 | 53 | 125 | 108 | 121 | 120 | 3 | 3 | 7 | 7 | 16 | 15 | 65 | 65 | 17 | 17 | 13 | 13 |
| V3-QM | 1500 | 1140 | 1193 | 849 | 27 | 20 | 66 | 46 | 177 | 123 | 624 | 445 | 127 | 86 | 172 | 129 | 307 | 291 | 15 | 15 | 12 | 12 | 71 | 63 | 128 | 122 | 52 | 50 | 29 | 29 |
| V3-HM | 1437 | 869 | 1227 | 699 | 14 | 13 | 26 | 22 | 547 | 116 | 400 | 335 | 89 | 79 | 151 | 134 | 210 | 170 | 11 | 11 | 2 | 2 | 81 | 46 | 69 | 65 | 26 | 25 | 21 | 21 |
| Total | 7579 | 5728 | 6285 | 4515 | 134 | 93 | 204 | 161 | 1234 | 626 | 3089 | 2317 | 662 | 520 | 962 | 796 | 1294 | 1214 | 61 | 59 | 49 | 44 | 282 | 234 | 570 | 553 | 208 | 201 | 124 | 123 |
| U/T | 5728/7579 | | 4515/6285 | | 93/134 | | 161/204 | | 626/1234 | | 2317/3089 | | 520/662 | | 796/962 | | 1214/1294 | | 59/61 | | 44/49 | | 234/282 | | 553/570 | | 201/208 | | 123/124 | |
| **IgG** | **Repertoire** | | **In-frame** | | **IGHJ1** | | **IGHJ2** | | **IGHJ3** | | **IGHJ4** | | **IGHJ5** | | **IGHJ6** | | **Out-of-frame** | | **IGHJ1** | | **IGHJ2** | | **IGHJ3** | | **IGHJ4** | | **IGHJ5** | | **IGHJ6** | |
| **Sample** | **total** | **unique** | **total** | **unique** | **total** | **unique** | **total** | **unique** | **total** | **unique** | **total** | **unique** | **total** | **unique** | **total** | **unique** | **total** | **unique** | **total** | **unique** | **total** | **unique** | **total** | **unique** | **total** | **unique** | **total** | **unique** | **total** | **unique** |
| V1-QG | 1249 | 736 | 998 | 510 | 17 | 14 | 20 | 15 | 94 | 66 | 630 | 272 | 118 | 66 | 119 | 77 | 251 | 226 | 5 | 5 | 6 | 6 | 23 | 22 | 122 | 112 | 75 | 62 | 20 | 19 |
| V1-HG | 1239 | 946 | 1007 | 728 | 18 | 17 | 33 | 25 | 70 | 57 | 584 | 403 | 146 | 107 | 156 | 119 | 232 | 218 | 12 | 12 | 11 | 11 | 26 | 25 | 105 | 95 | 49 | 46 | 29 | 29 |
| V2-QG | 418 | 305 | 342 | 235 | 9 | 9 | 5 | 4 | 30 | 28 | 202 | 120 | 43 | 32 | 53 | 42 | 76 | 70 | 3 | 3 | 0 | 0 | 9 | 9 | 42 | 38 | 12 | 12 | 10 | 8 |
| V2-HG | 771 | 534 | 646 | 414 | 12 | 10 | 23 | 15 | 89 | 52 | 318 | 218 | 95 | 57 | 109 | 62 | 125 | 120 | 4 | 4 | 1 | 1 | 23 | 21 | 63 | 61 | 24 | 23 | 10 | 10 |
| V3-QG | 662 | 390 | 512 | 252 | 10 | 7 | 25 | 13 | 52 | 37 | 346 | 138 | 36 | 25 | 43 | 32 | 150 | 138 | 7 | 7 | 10 | 10 | 23 | 22 | 81 | 71 | 18 | 18 | 11 | 10 |
| V3-HG | 1858 | 681 | 1549 | 443 | 7 | 5 | 21 | 11 | 106 | 47 | 1150 | 264 | 101 | 54 | 164 | 62 | 309 | 238 | 5 | 5 | 3 | 3 | 35 | 29 | 184 | 132 | 54 | 43 | 28 | 26 |
| Total | 6197 | 3592 | 5054 | 2582 | 73 | 62 | 127 | 83 | 441 | 287 | 3230 | 1415 | 539 | 341 | 644 | 394 | 1143 | 1010 | 36 | 36 | 31 | 31 | 139 | 128 | 597 | 509 | 232 | 204 | 108 | 102 |
| U/T | 3592/6197 | | 2582/5054 | | 62/37 | | 83/127 | | 287/441 | | 1415/3230 | | 341/539 | | 394/644 | | 1010/1143 | | 36/36 | | 32/32 | | 128/139 | | 109/597 | | 204/232 | | 102/108 | |

Note: U/T represents the ratio of unique to total sequences.

**Supplementary Table 5.** The sequence data of 6 IGHJ gene families in the in-frame and out-of-frame BCR-H repertoire from SLE volunteers

|  | **Repertoire** | | **In-frame** | | **IGHJ1** | | **IGHJ2** | | **IGHJ3** | | **IGHJ4** | | **IGHJ5** | | **IGHJ6** | |
| --- | --- | --- | --- | --- | --- | --- | --- | --- | --- | --- | --- | --- | --- | --- | --- | --- |
| **Sample** | **total** | **unique** | **total** | **unique** | **total** | **unique** | **total** | **unique** | **total** | **unique** | **total** | **unique** | **total** | **unique** | **total** | **unique** |
| S1-1 | 570721 | 16712 | 511377 | 14846 | 8109 | 336 | 24468 | 434 | 46695 | 1894 | 249093 | 7768 | 42177 | 1559 | 140835 | 2855 |
| S1-2 | 293305 | 7458 | 267141 | 6732 | 4740 | 165 | 14792 | 228 | 24525 | 811 | 127812 | 3510 | 24181 | 773 | 71091 | 1245 |
| S1-3 | 175479 | 4657 | 158713 | 4175 | 2976 | 105 | 8357 | 133 | 16310 | 527 | 78112 | 2102 | 14700 | 435 | 38258 | 874 |
| S2-1 | 547455 | 32930 | 482878 | 28856 | 7080 | 601 | 26120 | 907 | 54027 | 4321 | 207334 | 13558 | 41951 | 3018 | 146366 | 6452 |
| S2-2 | 614082 | 33199 | 540830 | 29027 | 7944 | 587 | 28538 | 941 | 62200 | 4451 | 234612 | 13873 | 45935 | 2993 | 161601 | 6182 |
| S2-3 | 542654 | 25268 | 479320 | 22069 | 7361 | 481 | 26428 | 727 | 49663 | 3243 | 208396 | 10594 | 42376 | 2372 | 145096 | 4653 |
| Total | 2201042 | 94956 | 1960939 | 83636 | 30849 | 1794 | 102275 | 2643 | 203757 | 12004 | 896963 | 40811 | 168944 | 8778 | 558151 | 17608 |
| U/T | 94956/2201042 | | 83636/1960939 | | 1794/30849 | | 1643/102275 | | 12004/203757 | | 40811/896963 | | 8778/168944 | | 17608/558151 | |
|  | **Repertoire** | | **Out-of-frame** | | **IGHJ1** | | **IGHJ2** | | **IGHJ3** | | **IGHJ4** | | **IGHJ5** | | **IGHJ6** | |
| **Sample** | **total** | **unique** | **total** | **unique** | **total** | **unique** | **total** | **unique** | **total** | **unique** | **total** | **unique** | **total** | **unique** | **total** | **unique** |
| S1-1 | 570721 | 16712 | 59344 | 1866 | 1490 | 62 | 2506 | 48 | 3218 | 131 | 25491 | 903 | 5718 | 227 | 20921 | 495 |
| S1-2 | 293305 | 7458 | 26164 | 726 | 790 | 24 | 955 | 19 | 1370 | 59 | 10597 | 341 | 2582 | 89 | 9870 | 194 |
| S1-3 | 175479 | 4657 | 16766 | 482 | 381 | 14 | 603 | 16 | 1738 | 47 | 7769 | 223 | 1213 | 58 | 5062 | 124 |
| S2-1 | 547455 | 32930 | 64577 | 4074 | 1515 | 119 | 2985 | 93 | 4769 | 384 | 28238 | 1981 | 6318 | 523 | 20752 | 974 |
| S2-2 | 614082 | 33199 | 73252 | 4172 | 1185 | 104 | 2394 | 91 | 5775 | 409 | 33766 | 2100 | 6563 | 494 | 23569 | 974 |
| S2-3 | 542654 | 25268 | 63334 | 3199 | 878 | 68 | 1892 | 65 | 4459 | 304 | 29474 | 1588 | 5430 | 390 | 21201 | 784 |
| Total | 2201042 | 94956 | 240103 | 11320 | 5361 | 323 | 9443 | 267 | 16870 | 1030 | 105861 | 5548 | 22394 | 1391 | 80174 | 2761 |
| U/T | 94956/2201042 | | 11320/240103 | | 323/5361 | | 267/9443 | | 1030/16870 | | 5548/105861 | | 1391/22394 | | 2761/80174 | |

Note: U/T represents the ratio of unique to total sequences.

**Supplementary Table 6.** The sequence data of 6 IGHJ gene families in the in-frame and out-of-frame BCR-H repertoire from breast cancer volunteers

|  | **Repertoire** | | **In-frame** | | **IGHJ1** | | **IGHJ2** | | **IGHJ3** | | **IGHJ4** | | | **IGHJ5** | | | | **IGHJ6** | |
| --- | --- | --- | --- | --- | --- | --- | --- | --- | --- | --- | --- | --- | --- | --- | --- | --- | --- | --- | --- |
| **Sample** | **total** | **unique** | **total** | **unique** | **total** | **unique** | **total** | **unique** | **total** | **unique** | **total** | | **unique** | **total** | **unique** | | **total** | | **unique** |
| B3-1 | 20155 | 2504 | 16665 | 2087 | 256 | 33 | 713 | 56 | 1400 | 208 | 8093 | | 1052 | 1474 | 208 | | 3985 | | 445 |
| B2-1 | 161983 | 19745 | 138798 | 16723 | 2001 | 241 | 6790 | 520 | 11738 | 1679 | 64510 | | 7828 | 12092 | 1689 | | 35977 | | 4066 |
| B1-1 | 23214 | 4559 | 20083 | 3902 | 248 | 54 | 613 | 89 | 1117 | 250 | 8867 | | 1703 | 2222 | 487 | | 6150 | | 1156 |
| B3-2 | 70031 | 7744 | 59316 | 6536 | 966 | 113 | 2555 | 196 | 4491 | 611 | 28080 | | 3181 | 4951 | 615 | | 15478 | | 1519 |
| B2-2 | 86602 | 8129 | 72850 | 6806 | 980 | 96 | 2896 | 178 | 6950 | 793 | 32509 | | 3096 | 6078 | 684 | | 21083 | | 1719 |
| B1-2 | 31795 | 5527 | 27230 | 4724 | 334 | 53 | 723 | 88 | 1369 | 284 | 12143 | | 2091 | 2917 | 582 | | 8362 | | 1386 |
| B3-3 | 145504 | 15503 | 122436 | 12971 | 1716 | 201 | 5579 | 375 | 9576 | 1262 | 56084 | | 6004 | 10698 | 1348 | | 33650 | | 3279 |
| B2-3 | 63404 | 9530 | 53985 | 8051 | 997 | 115 | 2309 | 231 | 4564 | 796 | 24077 | | 3636 | 5249 | 857 | | 14764 | | 2107 |
| B1-3 | 31184 | 4950 | 26916 | 4252 | 336 | 50 | 917 | 103 | 1349 | 275 | 11661 | | 1863 | 2927 | 513 | | 7925 | | 1176 |
| Total | 633872 | 78191 | 538279 | 66052 | 7834 | 956 | 23095 | 1836 | 42554 | 6158 | 246024 | | 30454 | 48608 | 6983 | | 147374 | | 16853 |
| U/T | 78191/633872 | | 66052/538279 | | 956/7834 | | 1836/23095 | | 6158/42554 | | 3054/246024 | | | 6983/48608 | | | | 16853/147374 | |
|  | **Repertoire** | | **Out-of-frame** | | **IGHJ1** | | **IGHJ2** | | **IGHJ3** | | **IGHJ4** | | | **IGHJ5** | | | | **IGHJ6** | |
| **Sample** | **total** | **unique** | **total** | **unique** | **total** | **unique** | **total** | **unique** | **total** | **unique** | **total** | **unique** | | **total** | | **unique** | | **total** | **unique** |
| B3-1 | 20155 | 2504 | 3490 | 417 | 54 | 7 | 194 | 12 | 224 | 38 | 1503 | 186 | | 408 | | 45 | | 870 | 106 |
| B2-1 | 161983 | 19745 | 23185 | 3022 | 405 | 54 | 658 | 50 | 1232 | 179 | 9976 | 1327 | | 2432 | | 373 | | 7114 | 853 |
| B1-1 | 23214 | 4559 | 3131 | 657 | 84 | 15 | 67 | 13 | 120 | 26 | 1269 | 250 | | 344 | | 83 | | 995 | 221 |
| B3-2 | 70031 | 7744 | 10715 | 1208 | 161 | 21 | 379 | 26 | 507 | 69 | 4919 | 556 | | 1136 | | 149 | | 2884 | 305 |
| B2-2 | 86602 | 8129 | 13752 | 1323 | 120 | 13 | 303 | 22 | 822 | 86 | 6037 | 586 | | 1476 | | 163 | | 4151 | 379 |
| B1-2 | 31795 | 5527 | 4565 | 803 | 143 | 19 | 184 | 14 | 200 | 44 | 1629 | 302 | | 476 | | 103 | | 1524 | 260 |
| B3-3 | 145504 | 15503 | 23068 | 2532 | 366 | 43 | 615 | 43 | 1139 | 137 | 10786 | 1187 | | 2742 | | 310 | | 5525 | 612 |
| B2-3 | 63404 | 9530 | 9419 | 1479 | 155 | 21 | 285 | 25 | 536 | 91 | 4136 | 645 | | 817 | | 159 | | 2858 | 456 |
| B1-3 | 31184 | 4950 | 4268 | 698 | 47 | 9 | 80 | 8 | 182 | 32 | 1600 | 247 | | 519 | | 102 | | 1573 | 253 |
| Total | 633872 | 78191 | 95593 | 12139 | 1535 | 202 | 2765 | 213 | 4962 | 702 | 41855 | 5286 | | 10350 | | 1487 | | 27494 | 3445 |
| U/T | 78191/633872 | | 12139/95593 | | 202/1535 | | 213/2765 | | 702/4962 | | 5286/41855 | | | 1487/10350 | | | | 3445/27494 | |

Note: U/T represents the ratio of unique to total sequences.

**Supplementary Table 7.** The sequence data of 6 IGHJ gene families in the in-frame and out-of-frame BCR-H repertoire from volunteers with a high titer of HbsAb

|  | **Repertoire** | | **In-frame** | | | **IGHJ1** | | | **IGHJ2** | | **IGHJ3** | | | **IGHJ4** | | | **IGHJ5** | | | **IGHJ6** | | |
| --- | --- | --- | --- | --- | --- | --- | --- | --- | --- | --- | --- | --- | --- | --- | --- | --- | --- | --- | --- | --- | --- | --- |
| **Sample** | **total** | **unique** | **total** | **unique** | | **total** | | **unique** | **total** | **unique** | **total** | | **unique** | **total** | | **unique** | **total** | **unique** | | **total** | | **unique** |
| HBsAg-1 | 763366 | 20379 | 369481 | 11070 | | 2109 | | 97 | 15192 | 297 | 19605 | | 993 | 133101 | | 4456 | 36486 | 1412 | | 152213 | | 3415 |
| HBsAg-2 | 889478 | 32277 | 409468 | 17329 | | 3473 | | 207 | 14856 | 376 | 21264 | | 1527 | 167115 | | 7928 | 34721 | 1869 | | 156580 | | 4843 |
| Total | 1652844 | 52656 | 778949 | 28399 | | 5582 | | 304 | 30048 | 673 | 40869 | | 2520 | 300216 | | 12384 | 71207 | 3281 | | 308793 | | 8258 |
| U/T | 52656/1652844 | | 28399/778949 | | | 304/5582 | | | 673/30048 | | 2520/40869 | | | 12384/300216 | | | 3281/71207 | | | 8258/308793 | | |
|  | **Repertoire** | | **Out-of-frame** | | | **IGHJ1** | | | **IGHJ2** | | **IGHJ3** | | | **IGHJ4** | | | **IGHJ5** | | | **IGHJ6** | | |
| **Sample** | **total** | **unique** | **total** | | **unique** | **total** | **unique** | | **total** | **unique** | **total** | **unique** | | **total** | **unique** | | **total** | | **unique** | **total** | **unique** | |
| HBsAg-1 | 763366 | 20379 | 393885 | | 9309 | 8015 | 242 | | 20156 | 228 | 15418 | 647 | | 154116 | 3686 | | 46467 | | 1497 | 131032 | 2577 | |
| HBsAg-2 | 889478 | 32277 | 480010 | | 14948 | 8564 | 392 | | 19155 | 355 | 19749 | 971 | | 173488 | 5859 | | 55282 | | 2335 | 176431 | 4222 | |
| Total | 1652844 | 52656 | 873895 | | 24257 | 16579 | 634 | | 39311 | 583 | 35167 | 1618 | | 327604 | 9545 | | 101749 | | 3832 | 307463 | 10006 | |
| U/T | 52656/1652844 | | 24257/873895 | | | 634/16579 | | | 583/39311 | | 1618/35167 | | | 9545/327604 | | | 3832/101749 | | | 6799/307463 | | |

Note: U/T represents the ratio of unique to total sequences.

**Supplementary Table 8.** The frequency of six IGHJ gene families in BCR-H repertoire from 19 publishing article

| **The Title of Paper** | **Object and number** | **Sample** | **BCR-H sequences (group)** | **The usage and distribution of IGHJ1&2&3&4&5&6** | **Sources of literature** |
| --- | --- | --- | --- | --- | --- |
| [1]Direct measurement of B-cell receptor repertoire's composition and variation in systemic lupus erythematosus (SLE) | 10 SLE patients  6 heathy controls | PBMC (DNA) | (1)SLE patients  (2)heathy controls | Figure 3 (C) (2 groups)  IGHJ4 >IGHJ6 > IGHJ5 > GHJ3 >IGHJ2 >IGHJ1 | [29] |
| [2]Clonal Characteristics of Circulating B Lymphocyte Repertoire in Primary Biliary Cholangitis（PBC) | 43 PBC patients  34 healthy volunteers | PBMC (RNA) | (1)PBC patients  (2)healthy volunteers | Figure 3 (B) (overlapping clones) (2 groups)  IGHJ4 > IGHJ3>IGHJ6 > GHJ5 >IGHJ2 >IGHJ1 | [30] |
| [3]Transitional B Cells in Early Human B Cell Development - Time to Revisit the Paradigm? | 19 healthy adult donors  (aged 24–86 years) | Sorting cells from Peripheral blood and Bone marrow | (1)Pre-B; (2)Immature B  (3)Transitional B ; (4)Naïve B | Figure 2 (A) (4 groups)  IGHJ4 >IGHJ6 >IGHJ3 >IGHJ5 >IGHJ2≈IGHJ1 | [31] |
| [4]High-Throughput Sequencing Reveals Immunological Characteristics of the TRB-/IgH-CDR3 Region of Umbilical Cord Blood | 20 healthy adults;  56 pregnant women  40 newborns | Umbilical Cord Blood  Peripheral blood | (1)Newborns; (2)Pregnant women; (3)adults | Figure 2 (F) (3 groups)  IGHJ4 >IGHJ6 >IGHJ3 >IGHJ5 >IGHJ2 >IGHJ1 | [32] |
| [5] Characterization of the B Cell Receptor Repertoire in the Intestinal Mucosa and of Tumor-Infiltrating Lymphocytes in Colorectal Adenoma and Carcinoma (CRC) | 6 healthy controls  4 AD patients  6 CRC.patients | Biopsies (RNA) | (1) healthy controls;(2)AD patients and CRC patients | Figure 6 (E) (2 groups)  Most of V pair to J were IGHJ4&IGHJ6 | [33] |
| [6]High-Throughput Single-Cell Analysis of B Cell Receptor Usage among Autoantigen-Specific Plasma Cells in Celiac Disease (CD) | 10 CD patients (Biopsies):  8 untreated consuming a normal diet; 2 treated consuming a gluten-free diet | Sorting Cells from Biopsies (RNA) | Celiac disease (CD) patients  (1)TG2+ ; (2) TG2- | Figure 1 (E) (2 groups)  IGHJ4 >IGHJ6 > IGHJ5 >GHJ3 >IGHJ2 >IGHJ1 | [34] |
| [7]The Human Thymus Is Enriched for Autoreactive B Cells | Thymus material was obtained from three children requiring surgery for congenital heart disease (2 months and 1.5 years of age); fetal BM samples was obtained from elective abortions (three donors); Pediatric BM samples were obtained from three children (aged 2–6years) [51] | Single-cell sorting from tissue (RNA) | (1)fetal BM; (2)pediatric BM;(3)pediatric thymus | Figure 2 (C) (3 groups)  Fetal BM:  IGHJ4 > IGHJ2> IGHJ3 > GHJ5>IGHJ6≈IGHJ1  pediatric BM:  IGHJ4 > IGHJ3 >IGHJ5 ≈IGHJ6≈IGHJ2 >IGHJ1  pediatric thymus:  IGHJ4 >IGHJ6 >IGHJ3 >IGHJ5 >IGHJ2 >IGHJ1 | [35] |
| [8]Unlike in Children with Allergic Asthma, IgE Transcripts from Preschool Children with Atopic Dermatitis Display Signs of Superantigen-Driven Activation | Five preschool children with atopic dermatitis | Peripheral blood (RNA) | IgE( IgM control ) :  (1)atopic dermatitis ; (2)allerjic asthma | Figure 1 (C) (2 groups)  IGHJ4 >IGHJ5 >IGHJ6 >IGHJ3 >IGHJ2 >IGHJ1 | [36] |
| [9]IMonitor: A Robust Pipeline for TCR and BCR Repertoire Analysis | Samples of peripheral blood from 2 healthy human donors (H-H-1, H-B-1) | Peripheral blood  (RNA) | 2 healthy donors | Figure 4 (k) (1 group)  IGHJ4 >IGHJ6 >IGHJ5 >IGHJ2 >IGHJ3> IGHJ1 | [37] |
| [10]The normal IGHV1-69-derived B-cell repertoire contains stereotypic patterns characteristic of unmutated CLL | 3 healthy persons: D1 (69 years); D2 (69 years) D3 (51 years); and age-matched to that of patients with CLL [52] | Peripheral blood  (RNA) | IGHJ use in normal B cells with IGHV1-69-DJ-C rearrangements:(1)previous study (n=26);(2)present study (n=72) | Figure 1. (2 groups)  IGHJ4 >IGHJ6 > IGHJ3> IGHJ5 >IGHJ2>IGHJ1 | [38] |
| [11]Antibody V(h) repertoire differences between resolving and chronically evolving hepatitis C virus infections | 7 healthy donors (HD); 6 patients (acute HCV infection, spontaneous resolvers, SR)  9 patients (chronic HCV infection, chronically evolving, CE) | Sorting cells (naive B cell clones and naive B cell clones) from Peripheral blood (DNA) | (1) healthy donors; (2) acute HCV infection, spontaneous resolvers; (3) chronic HCV infection, chronically evolving | Figure 1 (E F): (3 groups)  IGHJ4 >IGHJ6 > IGHJ3 >IGHJ5>IGHJ1  (No IGHJ2） | [39] |
| [12]Antibody repertoires in humanized NOD- scid- IL2Rgamma(null) mice and human B cells reveals human like diversification and tolerance checkpoints in the mouse | Two healthy females  humanized mouse spleens | human PBMCs; naive or total B cells from humanized mouse spleen; immature B cells pooled humanized mice immature B cells (RNA) | (1)Hu PBC-1; (2) Hu PBC-2 ; (3)HuMs-1NSpl; (4) HuMs- 2NSpl; (5) HuMs-3TSpl ; (6)HuMs-ImmB | Figure 2 (C) (6 groups)  IGHJ4 >IGHJ6 > GHJ3 >IGHJ5 >IGHJ2 >IGHJ1 | [40] |
| [13]Expressed antibody repertoires in human cord blood cells: 454 sequencing and IMGT/HighV-QUEST analysis of germline gene usage, junctional diversity, and somatic mutations | An African-American female baby  a Caucasian male baby | Two cord blood (RNA) | two babies IG:  (1) CB1 (productive);  (2) CB1 (unproductive);  (3) CB2 (productive);  (4) CB2 (unproductive) | Figure 1 (C) (4 groups)  IGHJ4 > IGHJ3 >IGHJ6 >IGHJ5 >IGHJ2 >IGHJ1 | [41] |
| [14]Tissue-specific expressed antibody variable gene repertoires | Healthy human subjects were obtained from a commercial source (Clontech). 39 peripheral leukocytes, 56 bone marrow, 15 small intestines, 13 lung, 7 stomach, 42 lymph node, 34 tonsil, 12 spleen and 25 thymuses | peripheral blood, bone marrow, mucosal tissues; lymph tissues (RNA) | (1) peripheral blood; (2) bone marrow; mucosal tissues (lung, small intestine, stomach); (3) lymphoid tissues (lymph node, tonsil, spleen and thymus). | Figure 1 (3 groups)  IGHJ4 >IGHJ6 > GHJ3 >IGHJ1>IGHJ2  (figure 1 No IGHJ5 ) | [42] |
| [15]Differences in the composition of the human antibody repertoire by B cell subsets in the blood | One healthy female subject (age 56) | Sorting cells (RNA) | (1)immature;(2)transitional;(3)Mature;(4)Memory;IgD+IgM; (5)memory IgD−IgM;(6) plasmacytes ;IgM;(7)Memory;IgD-IgG;(8)plasmacytes IgG | Figure 7 (8 groups）  IGHJ4 >IGHJ6 > IGHJ5 ≈ IGHJ3 >IGHJ2>IGHJ1 | [43] |
| [16]Prevalence and gene characteristics of antibodies with cofactor-induced HIV-1 specificity | Human immunodeficiency virus immune -globulin (HIVIg) was obtained through the NIH | The repertoire of human antibodies (gp120) | (1)Sensitive Abs  (2)Non-Sensitive Abs | Figure 6 (A [2]) (2 groups)  IGHJ4 >IGHJ6 > IGHJ3 ≈ IGHJ5 >IGHJ1>IGHJ2 | [44] |
| [17]Age-related aspects of human IgM+ B cell heterogeneity | Peripheral blood mononuclear cells were isolated from a total of 14 young (21–45 years) and 16 old (62–87 years) healthy volunteers. | Sorting cells from PBMCs（RNA） | (1)young naive;(2)old naive;(3)young IgM memory;(4)old IgM memory;(5)young IgM only CD27–;(6)old IgM only CD27–;(7)young IgM onlyCD27+;(8) old IgM onlyCD27+ | Figure 3 (A) (8 groups)  IGHJ4 >IGHJ6 >IGHJ3 ≈IGHJ5 >IGHJ2>IGHJ1 | [45] |
| [18]High-throughput sequencing of IgG B-cell receptors reveals frequent usage of the rearranged IGHV4-28/IGHJ4 gene in primary immune thrombocytopenia | Eleven adult chronic Primary immune thrombocytopenia patients and nine volunteer donors | PBMCs（RNA） | IgG-BCRs  (1)Primary immune thrombocytopenia; (2)volunteer donors | Figure 1 (2 groups)  IGHJ4 >IGHJ6 > IGHJ5 > IGHJ3 >IGHJ2>IGHJ1 | [46] |
| [19]IgM repertoire biodiversity is reduced in HIV-1 infection and systemic lupus erythematosus | Sixteen individuals: 4 healthy controls, 4 subjects with SLE, 4 therapy-naïve HIV, and 4 receiving combination antiretroviral therapy (cART) HIVTx | PBMCs（RNA） | IgM -BCRs  (454-deep pyrosequencing) | Figure 7 (C or D)  IGHJ4 >IGHJ3>IGHJ6>IGHJ5 >IGHJ2>IGHJ1 | [47] |

**Supplementary Table 9.** The composition and characteristics of human J-NONAMER--J-SPACER--J-HEPTAMER (9-23-7) recombination signal sequence (RSS) and J-REGION Subsequence & AA

| **Names** | **F** | **Accession**  **number** | **Location** | **J-NONAMER Subsequence** | **Location** | **J-SPACER**  **Subsequence** | **Location** | **J-HEPTAME Subsequence** | **J-REGION Location** | **J-REGION Subsequence & AA** |
| --- | --- | --- | --- | --- | --- | --- | --- | --- | --- | --- |
| IGHJ1*01 | **F** | [X97051](http://www.imgt.org/ligmdb/view.action?id=X97051) | 87524..87532 | ggtttctgt | 87533..87554 | agcccctggctcagggctgact [22]  (3a+4t+8c+7g) | 87555..87561 | caccgtg | 87562..87613 | gctgaatacttccagcactggggccagggcaccctggtcaccgtctcctcag  AEYFQHWGQGTLVTVSS [17] |
|  |  | [X86356](http://www.imgt.org/ligmdb/view.action?id=X86356) | 247..255 |  | 256..277 |  | 278..284 |  | 285..336 |  |
| IGHJ2*01 | **F** | [X97051](http://www.imgt.org/ligmdb/view.action?id=X97051) | 87731..87739 | tgtttttgt | 87740..87761 | atgggagaagcaggagggcaga [22]  (8a+1t+2c+11g) | 87762..87768 | ggctgtg | 87769..87821 | ctactggtacttcgatctctggggccgtggcaccctggtcactgtctcctcag  YWYFDLWGRGTLVTVSS [17] |
|  |  | [X86356](http://www.imgt.org/ligmdb/view.action?id=X86356) | 454..462 |  | 463..484 |  | 485..491 |  | 492-544 |  |
| IGHJ3*01 | **F** | [M25625](http://www.imgt.org/ligmdb/view.action?id=M25625) | 31..39 | ggtttgtgt | 40..62 | ctgggtctaggaacggactgtgt [23]  (4a+5t+4c+9g) | 63..69 | ccctgtg | 70..119 | tgatgcttttgatgtctggggccaagggacaatggtcaccgtctcttcag  DAFDVWGQGTMVTVSS [16] |
| IGHJ3*02 |  | [X97051](http://www.imgt.org/ligmdb/view.action?id=X97051) | 88345..88353 |  | 88354..88376 | ctgggcaggaacagggactgtgt [23]  (5a+4t+4c+10g) | 88377..88383 |  | 88384..88433 | tgatgcttttgatatctggggccaagggacaatggtcaccgtctcttcag  DAFDIWGQGTMVTVSS [16] |
|  |  | [X86356](http://www.imgt.org/ligmdb/view.action?id=X86356) | 1068..1076 |  | 1077..1099 |  | 1100..1106 |  | 1107..1156 |  |
| IGHJ4*01 | **F** | [J00256](http://www.imgt.org/ligmdb/view.action?id=J00256) | 1873..1881 | ggtttttgt | 1882..1904 | gcaccccttaatggggcctccca [23]  (4a+4t+10c+5g) | 1905..1911 | caatgtg | 1912..1959 | actactttgactactggggccaaggaaccctggtcaccgtctcctcag  YFDYWGQGTLVTVSS [15] |
| IGHJ4*02 |  | [X97051](http://www.imgt.org/ligmdb/view.action?id=X97051) | 88719..88727 |  | 88728..88750 |  | 88751..88757 |  | 88758..88805 |  |
|  |  | [X86356](http://www.imgt.org/ligmdb/view.action?id=X86356) | 1441..1449 |  | 1450..1472 |  | 1473..1479 |  | 1480..1527 |  |
| IGHJ4*03 |  | [M25625](http://www.imgt.org/ligmdb/view.action?id=M25625) | 407..415 |  | 416..438 |  | 439..445 |  | 446..493 |  |
| IGHJ5*01 | **F** | [M25625](http://www.imgt.org/ligmdb/view.action?id=M25625) | 855..863 | gttctttgt | 864..884 | cggggtctggcattgttgtca [21]  (2a+7t+4c+8g) | 885..891 | caatgtg | 892..942 | acaactggttcgactcctggggccaaggaaccctggtcaccgtctcctcag  NWFDPWGQGTLVTVSS [16] |
| IGHJ5*02 |  | [X97051](http://www.imgt.org/ligmdb/view.action?id=X97051) | 89118..89126 | gttcttgcc | 89127..89148 | tggggtcctggcattgttgtca [22]  (2a+8t+4c+8g) | 89149..89155 |  | 89156..89206 | acaactggttcgacccctggggccagggaaccctggtcaccgtctcctcag  NWFDSWGQGTLVTVSS [16] |
|  |  | [X86356](http://www.imgt.org/ligmdb/view.action?id=X86356) | 1840..1848 |  | 1849..1870 |  | 1871..1877 |  | 1878..1928 |  |
| IGHJ6*01 | **F** | [J00256](http://www.imgt.org/ligmdb/view.action?id=J00256) | 2909..2917 | ggtttttgt | 2918..2939 | ggggtgaggatggacattctgc [22]  (4a+5t+3c+10g) | 2940..2946 | cattgtg | 2947..3009 | attactactactactacggtatggacgtctgggggcaagggaccacggtcaccgtctcctcag YYYYYGMDVWGQGTTVTVSS [20] |
| IGHJ6*02 |  | [M25625](http://www.imgt.org/ligmdb/view.action?id=M25625) | 1442.22.1450 |  | 1451..1472 | tgggtgaggatggacattctgc [22]  (4a+6t+3c+9g) | 1473..1479 |  | 1480..1542 |  |
| IGHJ6*03 |  | [X97051](http://www.imgt.org/ligmdb/view.action?id=X97051) | 89722..89730 |  | 89731..89752 |  | 89753..89759 |  | 89760-89760..89822 |  |
|  |  | [X86356](http://www.imgt.org/ligmdb/view.action?id=X86356) | 2444..2452 |  | 2453..2474 | ggggtgaggatggacattctgc [22]  (4a+5t+3c+10g) | 2475..2481 |  | 2482..2543 | attactactactactacggtatggacgtctggggccaagggaccacggtcaccgtctcctca  YYYYYYMDVWGKGTTVTVSS [20] |
| IGHJ6*04 |  | [AJ879487](http://www.imgt.org/ligmdb/view.action?id=AJ879487) | 1..9 |  | 10..31 |  | 32..38 |  | 39..101 | attactactactactacggtatggacgtctggggcaaagggaccacggtcaccgtctcctcag  YYYYYGMDVWGKGTTVTVSS [20] |

**Supplementary Table 10.** The composition of human IGHD heptamer, spacer and nonamer (7-12-9 RSSs)

| **Accession number** | **Allele or Gene** | **Location** | **D-HEPTAMER** | **Location** | **D-SPACER Subsequence** | **Location** | **D-NONAMER** |
| --- | --- | --- | --- | --- | --- | --- | --- |
| [X97051](http://www.imgt.org/ligmdb/view.action?id=X97051) | IGHD1-1*01 | 33731..33737 | caccgtg | 33738..33749 | agaaaaactgtg | 33750..33758 | tccaaaact |
| [X97051](http://www.imgt.org/ligmdb/view.action?id=X97051) | IGHD1-7*01 | 43317..43323 | cactgtg | 43324..43335 | agaaaagcttcg | 43336..43344 | tccaaaacg |
| [X97051](http://www.imgt.org/ligmdb/view.action?id=X97051) | IGHD1-14*01 | 52585..52591 | cactgtc | 52592..52603 | agaatagctacg | 52604..52612 | tcaaaaact |
| [X97051](http://www.imgt.org/ligmdb/view.action?id=X97051) | IGHD1-20*01 | 62032..62038 | caccgtg | 62039..62050 | agaaaaactgtg | 62051..62059 | tccaaaact |
| [X97051](http://www.imgt.org/ligmdb/view.action?id=X97051) | IGHD1-26*01 | 72189..72195 | cactgtg | 72196..72207 | agaaaagctatg | 72208..72216 | tccaaaact |
| [X97051](http://www.imgt.org/ligmdb/view.action?id=X97051) | IGHD2-2*02 | 36398..36404 | cacagtg | 36405..36416 | acacagccccat | 36417..36425 | tcccaaagc |
| [X97051](http://www.imgt.org/ligmdb/view.action?id=X97051) | IGHD2-8*01 | 46013..46019 | cacagtg | 46020..46031 | acacagccccat | 46032..46040 | tcccaaagc |
| [X97051](http://www.imgt.org/ligmdb/view.action?id=X97051) | IGHD2-15*01 | 55266..55272 | cacagtg | 55273..55284 | acacagacccat | 55285..55293 | tcccaaagc |
| [X97051](http://www.imgt.org/ligmdb/view.action?id=X97051) | IGHD2-21*02 | 64672..64678 | cacagtg | 64679..64690 | acacaaccccat | 64691..64699 | tcctaaagc |
| [X97051](http://www.imgt.org/ligmdb/view.action?id=X97051) | IGHD3-3*01 | 38865..38871 | cacagtg | 38872..38883 | tcacagagtcca | 38884..38892 | tcaaaaacc |
| [X97051](http://www.imgt.org/ligmdb/view.action?id=X97051) | IGHD3-9*01 | 48543..48549 | cacagtg | 48550..48561 | tcacagagtcca | 48562..48570 | tcaaaaacc |
| [X97051](http://www.imgt.org/ligmdb/view.action?id=X97051) | IGHD3-10*01 | 48727..48733 | cacagtg | 48734..48745 | tcacagagtcca | 48746..48754 | tcaaaaacc |
| [X97051](http://www.imgt.org/ligmdb/view.action?id=X97051) | IGHD3-16*02 | 57589..57595 | cacagca | 57596..57607 | tcacacggtcca | 57608..57616 | tcagaaacc |
| [X97051](http://www.imgt.org/ligmdb/view.action?id=X97051) | IGHD3-22*01 | 67192..67198 | cacagtg | 67199..67210 | tcacagagtcca | 67211..67219 | tcaaaaact |
| [X97051](http://www.imgt.org/ligmdb/view.action?id=X97051) | IGHD4-4*01 | 40002..40008 | cacagtg | 40009..40020 | atgaacccagca | 40021..40029 | gcaaaaact |
| [X97051](http://www.imgt.org/ligmdb/view.action?id=X97051) | IGHD4-11*01 | 49607..49613 | catagtg | 49614..49625 | atgaacccagtg | 49626..49634 | gcaaaaact |
| [X97051](http://www.imgt.org/ligmdb/view.action?id=X97051) | IGHD4-17*01 | 58715..58721 | cacagtg | 58722..58733 | atgaaactagca | 58734..58742 | gcaaaaact |
| [X97051](http://www.imgt.org/ligmdb/view.action?id=X97051) | IGHD4-23*01 | 68353..68359 | cacagtg | 68360..68371 | atgaaaccagca | 68372..68380 | gcaaaaact |
| [X97051](http://www.imgt.org/ligmdb/view.action?id=X97051) | IGHD5-5*01 | 40967..40973 | cacagtg | 40974..40985 | gtgctgcccata | 40986..40994 | gcagcaacc |
| [X97051](http://www.imgt.org/ligmdb/view.action?id=X97051) | IGHD5-12*01 | 50574..50580 | cacagtg | 50581..50592 | gtgccgcccata | 50593..50601 | gcagcaacc |
| [X97051](http://www.imgt.org/ligmdb/view.action?id=X97051) | IGHD5-18*01 | 59681..59687 | cacagtg | 59688..59699 | gtgctgcccata | 59700..59708 | gcagcaacc |
| [X97051](http://www.imgt.org/ligmdb/view.action?id=X97051) | IGHD5-24*01 | 69320..69326 | cacagtg | 69327..69338 | gtgccgcccata | 69339..69347 | gcagcaacc |
| [X97051](http://www.imgt.org/ligmdb/view.action?id=X97051) | IGHD6-6*01 | 42813..42819 | cacagtg | 42820..42831 | acactcgccagg | 42832..42840 | ccagaaacc |
| [X97051](http://www.imgt.org/ligmdb/view.action?id=X97051) | IGHD6-13*01 | 52081..52087 | cacagtg | 52088..52099 | acactcacccag | 52100..52108 | ccagaaacc |
| [X97051](http://www.imgt.org/ligmdb/view.action?id=X97051) | IGHD6-19*01 | 61524..61530 | cacagtg | 61531..61542 | acactcgccagg | 61543..61551 | ccagaaacc |
| [X97051](http://www.imgt.org/ligmdb/view.action?id=X97051) | IGHD6-25*01 | 71684..71690 | cacaatg | 71691..71702 | acactgggcagg | 71703..71711 | acagaaacc |
| [X97051](http://www.imgt.org/ligmdb/view.action?id=X97051) | IGHD7-27*01 | 87470..87476 | cacagtg | 87477..87488 | attggcagctct | 87489..87497 | acaaaaacc |

**Supplementary Table 11.** The pairing of human IGHD heptamer-spacer-nonamer (7-12-9）RSSs and IGHJ nonamer-Spacer-heptamer (9-23-7) RSSs

| **Allele or Gene** | **D-HEPTAMER** | **IGHJ1(**caccgtg) **to D-HEPTAMER** | **IGHJ2 (**ggctgtg)**to D-HEPTAMER** | **IGHJ3 (**ccctgtg) **to D-HEPTAMER** | **IGHJ6(**cattgtg) **to D-HEPTAMER** | **IGHJ4&IGH5 (**caatgtg) **to D-HEPTAMER** | **D-NONAMER** | **IGHJ1(**Ggtttctgt) **to D-NONAMER** | **IGHJ2(**tgtttttgt)) **to D-NONAMER** | **IGHJ3(**Ggtttgtgt)) **to D-NONAMER** | **IGHJ5(**gttctttgt)) **to D-NONAMER** | **IGHJ4&IGHJ6**  **-(**Ggtttttgt)) **to D-NONAMER** |
| --- | --- | --- | --- | --- | --- | --- | --- | --- | --- | --- | --- | --- |
| IGHD1-1*01 | Caccgtg | caccgtg | caccgtg | caccgtg | caccgtg | caccgtg | tccaaaact | tccaaaact | tccaaaact | tccaaaact | tccaaaact | tccaaaact |
| IGHD1-7*01 | cactgtg | cactgtg | cactgtg | cactgtg | cactgtg | cactgtg | tccaaaacg | tccaaaacg | tccaaaacg | tccaaaacg | tccaaaacg | tccaaaacg |
| IGHD1-14*01 | cactgtc | cactgtc | cactgtc | cactgtc | cactgtc | cactgtc | tcaaaaact | tcaaaaact | tcaaaaact | tcaaaaact | tcaaaaact | tcaaaaact |
| IGHD1-20*01 | caccgtg | caccgtg | caccgtg | caccgtg | caccgtg | caccgtg | tccaaaact | tccaaaact | tccaaaact | tccaaaact | tccaaaact | tccaaaact |
| IGHD1-26*01 | cactgtg | cactgtg | cactgtg | cactgtg | cactgtg | cactgtg | tccaaaact | tccaaaact | tccaaaact | tccaaaact | tccaaaact | tccaaaact |
| IGHD2-2*02 | cacagtg | cacagtg | cacagtg | cacagtg | cacagtg | cacagtg | tcccaaagc | tcccaaagc | tcccaaagc | tcccaaagc | tcccaaagc | tcccaaagc |
| IGHD2-8*01 | cacagtg | cacagtg | cacagtg | cacagtg | cacagtg | cacagtg | tcccaaagc | tcccaaagc | tcccaaagc | tcccaaagc | tcccaaagc | tcccaaagc |
| IGHD2-15*01 | cacagtg | cacagtg | cacagtg | cacagtg | cacagtg | cacagtg | tcccaaagc | tcccaaagc | tcccaaagc | tcccaaagc | tcccaaagc | tcccaaagc |
| IGHD2-21*02 | cacagtg | cacagtg | cacagtg | cacagtg | cacagtg | cacagtg | tcctaaagc | tcctaaagc | tcctaaagc | tcctaaagc | tcctaaagc | tcctaaagc |
| IGHD3-3*01 | cacagtg | cacagtg | cacagtg | cacagtg | cacagtg | cacagtg | tcaaaaacc | tcaaaaacc | tcaaaaacc | tcaaaaacc | tcaaaaacc | tcaaaaacc |
| IGHD3-9*01 | cacagtg | cacagtg | cacagtg | cacagtg | cacagtg | cacagtg | tcaaaaacc | tcaaaaacc | tcaaaaacc | tcaaaaacc | tcaaaaacc | tcaaaaacc |
| IGHD3-10*01 | cacagtg | cacagtg | cacagtg | cacagtg | cacagtg | cacagtg | tcaaaaacc | tcaaaaacc | tcaaaaacc | tcaaaaacc | tcaaaaacc | tcaaaaacc |
| IGHD3-16*02 | cacagca | cacagca | cacagca | cacagca | cacagca | cacagca | tcagaaacc | tcagaaacc | tcagaaacc | tcagaaacc | tcagaaacc | tcagaaacc |
| IGHD3-22*01 | cacagtg | cacagtg | cacagtg | cacagtg | cacagtg | cacagtg | tcaaaaact | tcaaaaact | tcaaaaact | tcaaaaact | tcaaaaact | tcaaaaact |
| IGHD4-4*01 | cacagtg | cacagtg | cacagtg | cacagtg | cacagtg | cacagtg | gcaaaaact | gcaaaaact | gcaaaaact | gcaaaaact | gcaaaaact | gcaaaaact |
| IGHD4-11*01 | catagtg | catagtg | catagtg | catagtg | catagtg | catagtg | gcaaaaact | gcaaaaact | gcaaaaact | gcaaaaact | gcaaaaact | gcaaaaact |
| IGHD4-17*01 | cacagtg | cacagtg | cacagtg | cacagtg | cacagtg | cacagtg | gcaaaaact | gcaaaaact | gcaaaaact | gcaaaaact | gcaaaaact | gcaaaaact |
| IGHD4-23*01 | cacagtg | cacagtg | cacagtg | cacagtg | cacagtg | cacagtg | gcaaaaact | gcaaaaact | gcaaaaact | gcaaaaact | gcaaaaact | gcaaaaact |
| IGHD5-5*01 | cacagtg | cacagtg | cacagtg | cacagtg | cacagtg | cacagtg | gcagcaacc | gcagcaacc | gcagcaacc | gcagcaacc | gcagcaacc | gcagcaacc |
| IGHD5-12*01 | cacagtg | cacagtg | cacagtg | cacagtg | cacagtg | cacagtg | gcagcaacc | gcagcaacc | gcagcaacc | gcagcaacc | gcagcaacc | gcagcaacc |
| IGHD5-18*01 | cacagtg | cacagtg | cacagtg | cacagtg | cacagtg | cacagtg | gcagcaacc | gcagcaacc | gcagcaacc | gcagcaacc | gcagcaacc | gcagcaacc |
| IGHD5-24*01 | cacagtg | cacagtg | cacagtg | cacagtg | cacagtg | cacagtg | gcagcaacc | gcagcaacc | gcagcaacc | gcagcaacc | gcagcaacc | gcagcaacc |
| IGHD6-6*01 | cacagtg | cacagtg | cacagtg | cacagtg | cacagtg | cacagtg | ccagaaacc | ccagaaacc | ccagaaacc | ccagaaacc | ccagaaacc | ccagaaacc |
| IGHD6-13*01 | cacagtg | cacagtg | cacagtg | cacagtg | cacagtg | cacagtg | ccagaaacc | ccagaaacc | ccagaaacc | ccagaaacc | ccagaaacc | ccagaaacc |
| IGHD6-19*01 | cacagtg | cacagtg | cacagtg | cacagtg | cacagtg | cacagtg | ccagaaacc | ccagaaacc | ccagaaacc | ccagaaacc | ccagaaacc | ccagaaacc |
| IGHD6-25*01 | cacaatg | cacaatg | cacaatg | cacaatg | cacaatg | cacaatg | acagaaacc | acagaaacc | acagaaacc | acagaaacc | acagaaacc | acagaaacc |
| IGHD7-27*01 | cacagtg | cacagtg | cacagtg | cacagtg | cacagtg | cacagtg | acaaaaacc | acaaaaacc | acaaaaacc | acaaaaacc | acaaaaacc | acaaaaacc |
|  |  | 32/189 | 60/189 | 35/189 | 35/189 | 36/189 |  | 69/243 | 82/243 | 76/243 | 105/243 | 64/243 |
